# STASCAN deciphers fine-resolution cell-distribution maps in spatial transcriptomics by deep learning

**DOI:** 10.1101/2023.09.02.556029

**Authors:** Ying Wu, Jia-Yi Zhou, Bofei Yao, Guanshen Cui, Yong-Liang Zhao, Chun-Chun Gao, Ying Yang, Shihua Zhang, Yun-Gui Yang

**Affiliations:** CAS Key Laboratory of Genomic and Precision Medicine, Beijing Institute of Genomics, Chinese Academy of Sciences and China National Center for Bioinformation, Beijing 100101, China; University of Chinese Academy of Sciences, Beijing 100049, China; College of Future Technology, University of Chinese Academy of Sciences, Beijing 100049, China; NCMIS, CEMS, RCSDS, Academy of Mathematics and Systems Science, Chinese Academy of Sciences, Beijing 100190, China; School of Mathematical Sciences, University of Chinese Academy of Sciences, Beijing 100049, China; Key Laboratory of Systems Health Science of Zhejiang Province, School of Life Science, Hangzhou Institute for Advanced Study, University of Chinese Academy of Sciences, Hangzhou, 310024, China; Sino-Danish College, University of Chinese Academy of Sciences, Beijing 101408, China; Institute of Stem Cell and Regeneration, Chinese Academy of Sciences, Beijing 100101, China

**Keywords:** Spatial transcriptomics, Cell annotation, Deep learning, Imputation, Multimodal data integration

## Abstract

**Background:** The spatial transcriptomics (ST) technologies have been widely applied to decode the spatial distribution of cells by resolving gene expression profiles in tissues. However, a fine-resolved spatial cell map is still limited by algorithmic tools and sequencing techniques.

**Results:** Here we develop a novel deep learning approach, STASCAN, which could define the spatial cellular distribution of both captured and uncharted areas by cell feature learning that combines gene expression profiles and histology images. STASCAN additionally adopts optional transfer learning and pseudo-labeling methods to improve the accuracy of the cell-type prediction from images. We have successfully applied STASCAN to enhance cell resolution, and revealed finer organizational structures across diverse datasets from various species and tissues generated from 10× Visium technology. STASCAN improves cell resolution of *Schmidtea mediterranea* datasets by six times and reconstructs more detailed 3D cell-type models. Furthermore, STASCAN could accurately pinpoint the boundaries of distinct cell layers in human intestinal tissue, specifically identify a micrometer-scale smooth muscle bundle structure in consistent with anatomic insights in human lung tissue, and redraw the spatial structural variation with enhanced cell patterns in human myocardial infarction tissue. Additionally, through STASCAN on embryonic mouse brain datasets generated by DBiT-derived MISAR-seq technology, the increased cellular resolution and distinct anatomical tissue domains with cell-type niches are revealed. Collectively, STASCAN is compatible with different ST technologies and has notable advantages in generating cell maps solely from histology images, thereby enhancing the spatial cellular resolution.

**Conclusions:** In short, STASCAN displays significant advantages in deciphering higher-resolution cellular distribution, resolving enhanced organizational structures and demonstrating its potential applications in exploring cell-cell interactions within the tissue microenvironment.

## Introduction

The spatial distribution and function of cells are intimately related, and their characterization can provide valuable insights into their function and potential impact on biological processes, including development and disease^1^. The emerging spatial transcriptomics (ST) technologies allow to capture gene expression while preserving spatial context in tissue, which improves our understanding of the structure and the cellular composition of different organs across different species^2^.

Current ST technologies are typically divided into two categories: (1) imaging-based approaches that capture genes corresponding to the targeted probes through *in situ* sequencing or *in situ* hybridization, which can achieve single-cell resolution but are limited by low throughput, biased coverage for pre-selected gene targets, and expensive specialized equipment; and (2) next-generation sequencing (NGS)-based approaches that capture transcripts from tissues combined with encoded spatial positional information before sequencing^3^. By contrast, NGS-based approaches have been more widely used due to their advantages of higher throughput, unbiased coverage for transcriptome, and more accessible commercial products^4^.

However, NGS-based approaches still suffer from several limitations. The significant one is the spatial resolution which is restricted by the area and sparsity of captured domains (defined as spots)^5^. For example, the spots of 10× Visium are 55 μm in diameter with a space of 100 μm, which are too large to capture the cells at single-cell resolution and too sparse to measure cells in the uncharted areas between the captured domains^6^. In addition, DBiT-seq^7^ was designed to produce smaller and more densely spots (10, 25, or 50 μm in width). Even though it provides higher spatial resolution, the transcriptional information in the uncharted areas is still inevitably lost. Other approaches, such as Slide-seq^8^ and Stereo-seq^9^, also aim to achieve single-cell or subcellular resolution by using small and densely packed spots. However, they are also affected by the fact of multiple fractions of cells within a single spot.

Moreover, the uncaptured cells in uncharted areas lead to a limited spatial cellular resolution in not only 2D but also 3D levels. Currently, ST technologies enable us to depict cellular maps from a regional tissue section. By continuously sectioning the tissue, 3D cellular maps of organs can be constructed. However, the high-cost only allows a small portion of consecutive tissue sections be sequenced, incurring the problem of uncharted areas along the z-axis and ultimate low 3D resolution ^10^.

To improve the cellular resolution of ST technologies, current computational methods typically resolve gene profiles to perform cell-type deconvolution by integrating ST data with signatures of single-cell references and annotating cell types for captured domains^11^, such as Cell2location^12^, Seurat^13^, and RCTD^14^. However, the deconvolution methods are easily affected by the potential ‘dropout’ of marker genes in ST data and the inaccuracies in single-cell reference data^15^. More importantly, these methods only aim to improve computationally cellular resolution within captured domains by inferring the proportion or abundance of cell type. But how to enhance the spatial cellular resolution by predicting cell distribution in uncharted areas and imputing cell distribution between tissue sections in the z-axis remains to be solved.

In addition to the gene expression information derived from sequencing data, morphological information is also usually used to identify and characterize cell types of medical images^16^. Current deconvolution methods tend to focus on gene expression data, often overlooking the morphological information carried by images of ST datasets, resulting in a waste of image resources. More importantly, morphological information can also contribute to increase the accuracy of cell annotation. Some emerging computational methods integrating gene expression and morphological information have been developed for ST data. For instance, Tangram^17^ synthesizes histological images to estimate the proportion of cells in each spot during single-cell deconvolution, and MUSE^18^ combines transcriptional profiles and morphological features to characterize the cells and tissue regions. However, none of them involve the improvment of spatial cellular resolution in uncharted areas. Furthermore, some developed methods leverage ST data and histology images to enhance spatial gene profiles in captured domains and even uncharted areas^19^. Nevertheless, due to the limitation of cell-type deconvolution methods mentioned above and the need for refining gene expression-imputing models^20^, there are still obstacles to directly use the enhanced gene expression profile to predict cell types.

To address this issue, we introduce STASCAN, a Spatial TrAnscriptomics-driven Spatially Cellular ANnotation tool. STASCAN enables cell-type predictions in uncharted areas across tissue sections and subdivided-resolved annotation of cells within captured areas, thereby greatly enhancing spatial cellular resolution. In addition, STASCAN succeeds in generating cell distribution maps solely from histological images of adjacent sections, enabling the construction of a more detailed 3D cellular atlas of organs with reduced experimental costs. Furthermore, we evaluated the applicability of STASCAN in diverse datasets from different ST technologies and consistently observed significant improvements of STASCAN in cell granularity and comprehensive characterization of resolved cell patterns. For instance, STASCAN identified a micrometer-scale oval structure, confirmed as a smooth muscle bundle near the tracheal wall, which has not been identified by other methods. Moreover, STASCAN provided a refined distribution of cell-type niches in human cardiac tissue and mouse embryonic tissue, facilitating the understanding of disease and development.

## Results

### Overview of STASCAN

STASCAN adopts a deep learning model to utilize both the spatial gene expression profiles from the ST technology and the corresponding histological images, to depict a fine-resolution cell distribution map in tissues (**Fig. 1a**). Firstly, STASCAN extracts the spot images from slide images based on location information, and infers highly reliable cell labels for each spot based on spatial gene expression using deconvolution during the pre-labeling process. Secondly, STASCAN constructs a base convolutional neural network (CNN) model and trains the base CNN model using the cell-type labeled spot images as input. Additionally, STASCAN provides optional section-specific training, which can fine-tune the base CNN model through transfer learning to improve the prediction accuracy for a specific single section. Finally, through ample training, STASCAN can accurately predict cell types solely based on histological images (**Fig. 1a**, **Supplementary Fig.1** and **Methods**).

**Fig. 1.**
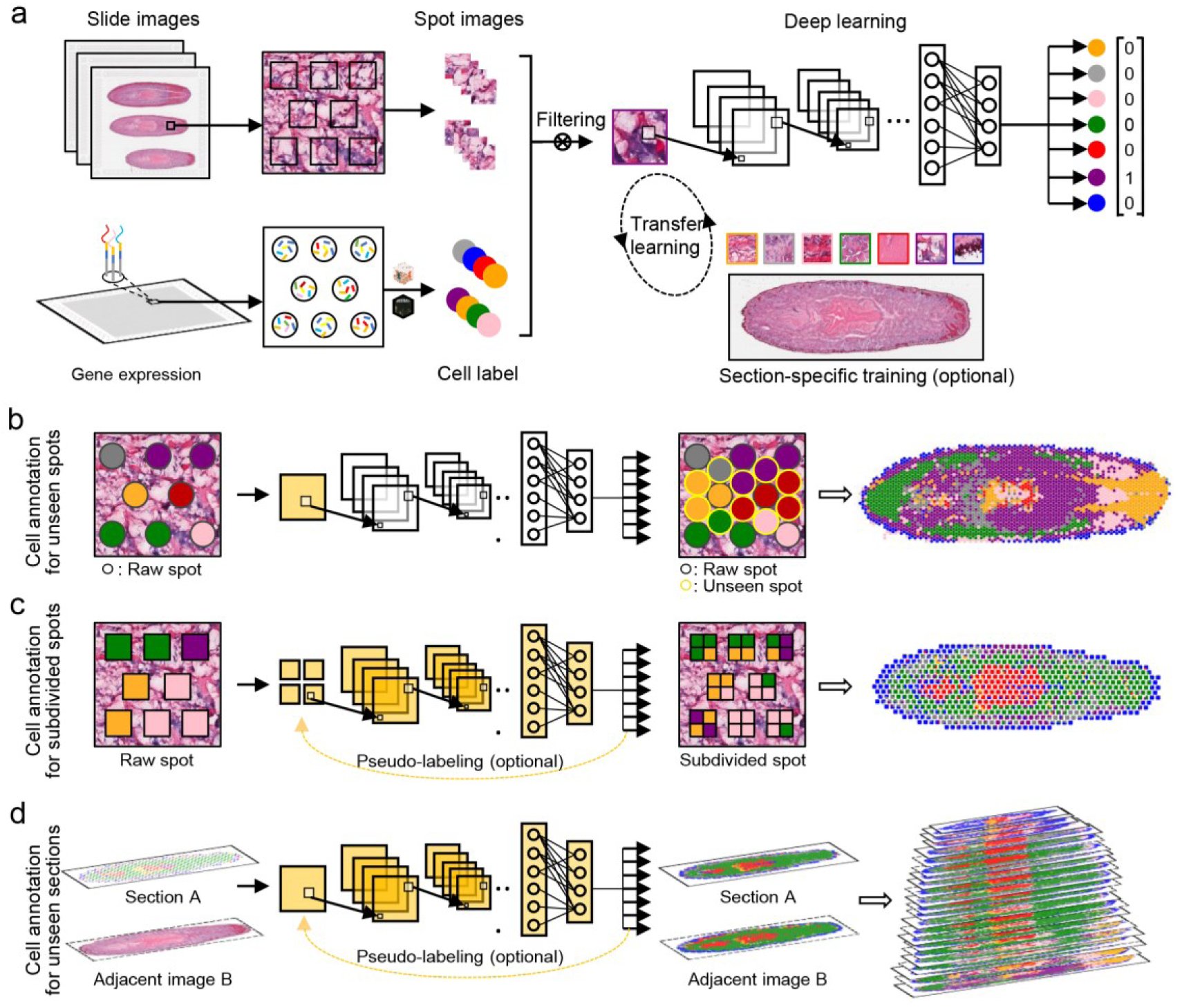
Overview of STASCAN. **(a)** Architecture of the STASCAN framework. STASCAN infers fine-resolution cell distribution with spatial transcriptomic (ST) data based on a deep learning model. Preprocessed spot images and dominant cell labels from slide images and gene expression data are taken as input to train a base CNN model. STASCAN provides an optional process to conduct section-specific training, which performs transfer learning to learn the characteristics of specific sections. **(b)** Architecture of cell annotation for unseen spots. STASCAN infers cell types for unseen spot images to decipher the super-resolution cell distribution of a ST slide. Unseen spots, the artificial spots among the unmeasured area. **(c)** Architecture of cell annotation for subdivided spots. STASCAN infers cell types of subdivided spot images to decipher the sub-resolution cell distribution of a ST slide. Raw spots, spots at raw resolution. Subdivided spots, spots by dividing raw spots for high resolution. **(d)** Architecture of cell annotation for unseen sections. STASCAN predicts cell types of simulated spots on adjacent images by learning characteristics of prior spot images from ST sections, depicting 3D cell distribution of tissue. Sections, ST sections with sequencing data. Adjacent images, adjacent H&E images without sequencing data.

STASCAN is further designed into three application modules: (1) cell annotation for the embedded unseen spots in uncharted areas, which is based on learning image features from the measured raw spots, assigning predicted cell type for each unseen spot, and merging the unseen and raw spots to achieve a super-resolved cell distribution (**Fig. 1b**); (2) cell annotation for subdivided spots, utilizing features learned from sub-spot images with optional pseudo-labels to obtain a sub-resolution cell distribution (**Fig. 1c**); (3) cell annotation for unseen sections, which learns the spot images from measured ST sections with optional pseudo-labels to predict the cell distribution on adjacent uncharted section images from the consecutive sections for constructing 3D cell models (**Fig. 1d**).

### STASCAN enables more precise cell annotation and cell types prediction solely from images

To quantitatively evaluate the performance of STASCAN, we initially applied it in a comprehensive planarian (*Schmidtea mediterranea*) dataset generated by 10× Visium technology, which includes ten sequenced ST sections (containing both spatial gene expression data and histological images) and nine unsequenced sections adjacent to ST sections (only containing histological images)^21^. Given the comprehensiveness of the planarian dataset, we first constructed a base model using 1,829 spot images extracted from 10 collected sections to learn the features of seven main cell types identified by seuqnecing information, including epidermal, gut, muscle, neoblast, neuronal, parenchymal, and secretory cells (**Supplementary Fig. 2a, b**, **Supplementary Table 1** and **Methods**). Although there was uncertainty in predicting neoblast and parenchymal cell types due to the scarcity of training samples of these two, most cell types were annotated with a recall rate of over 78% (**Fig. 2a** and **Supplementary Fig. 2c**). In addition, the learned model showed excellent accuracy in predicting cell types, with the area under the curve (AUC) calculated from the receiver operating characteristic (ROC) curves reaching as high as 0.936 to 0.996 (**Fig. 2b**). Besides, considering the potential batch effect among different ST sections, we performed section-specific training based on the base model (**Methods**). The results from the section-specific model showed a significant improvement in accuracy with higher AUC values compared with the base model, indicating that section-specific training is beneficial for the improved prediction performance of the whole frame (**Fig. 2c**).

**Fig. 2.**
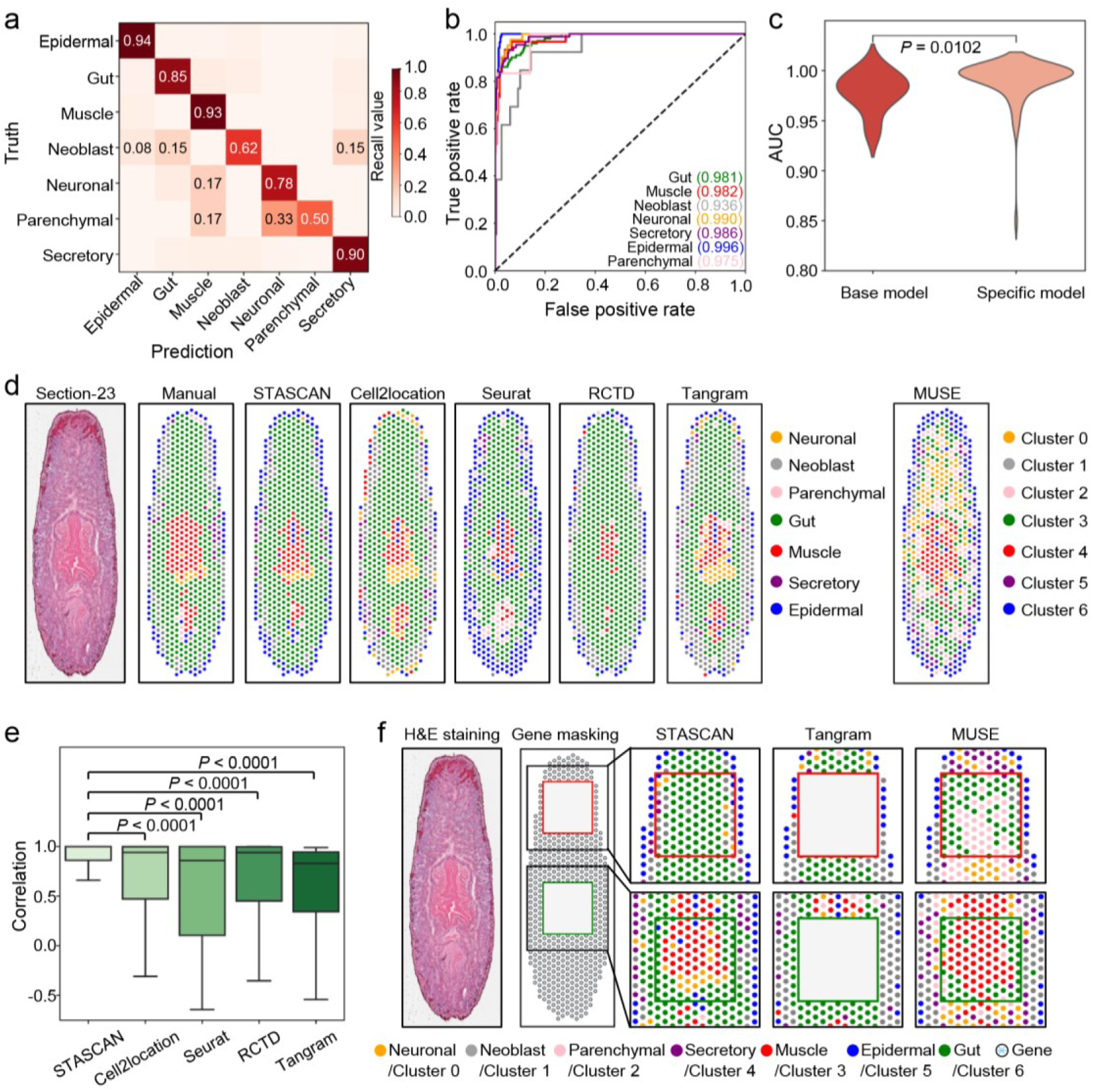
Evaluation of STASCAN in 10× Visium planarian dataset. **(a)** Heatmap showing the confusion matrix of the prediction and truth value of the base model for the 5-fold cross-validation in the planarian dataset. The number in each box represents the recall value. **(b)** ROC curves of each major cell type of the base model in the planarian dataset. The number in each pair of parentheses represents the AUC value. **(c)** Violin plot showing the AUC values of the base model (n = 7) and specific model (n = 70), respectively. P value is calculated using two-sided Wilcoxon signed-rank tests. **(d)** Cellular distribution of section 23 in the planarian dataset. Left, H&E staining image showing general morphology and spatial plots (x and y axis) depicting the distribution of the major cell types annotated by manual annotation, STASCAN, Cell2location, Seurat, RCTD, and Tangram. Right, spatial plots (x and y axis) depicting the distribution of cell clusters assigned by MUSE. **(e)** Boxplot showing the Pearson correlation between cell distribution of section 23 annotated by manual and alternative methods, respectively (n = 683, 683, 683, 681, 683). *P* values are calculated using two-sided Wilcoxon signed-rank tests. **(f)** Cellular distribution of section 23 in the planarian dataset with partly gene expression masking. Left, H&E staining image showing general morphology. Middle, schematic of gene expression masking. Right, zoomed regions of spatial plots (x and y axis) depicting the cell distribution predicted by STASCAN, Tangram, and MUSE.

To compare the performance of STASCAN in predicting dominant cell types on raw spots with other methods, we similarly performed Cell2location^12^, Seurat^13^, and RCTD^14^ on the planarian dataset. We initially annotated the cell types of each raw spot by manual according to the morphologic features of corresponding spot images, which considered as the ground truth. The cell distribution predicted by STASCAN is highly correlated with that of manual annotations, significantly superior to other methods (**Fig. 2d, e** and **Methods**). We also observed that alternative methods result in varied degrees of biases for cell annotation. For example, Cell2location could characterize most cell distribution but with a low sensitivity for epidermal cells; Seurat showed strong annotation bias for epidermal cells, leading to mislabeling for other cell types; and RCTD displayed some positive annotation but lost annotation information for most neuronal and secretory cells.

We also compared STASCAN with other methods that utilize both morphological and transcriptional features for ST data analysis. Tangram^17^ effectively illustrated the distribution of the majority of cells, but exhibited a slight bias towards neuronal cells and a reduced sensitivity in detecting epidermal cells (**Fig. 2d** and **e**). On the other hand, MUSE^18^ characterized tissue regions by identifying spot clusters, yet these clusters appeared relatively scattered, and failed to represent corresponding structures in the planarian dataset (**Fig. 2d**). In contrast, our STASCAN displayed more precise performance in prediction, with the capability of accurately pinpointing the spatial distribution of seven main types of cells, in accord with their known biological functions^22^ (**Fig. 2d**, and **Supplementary Fig. 2d,e**). For example, corresponding to clear tissue structures visible through hematoxylin and eosin (H&E) staining, the epidermal cells draw the contour of the planarian body, gut cells mark the location of the intestine, and muscle cells along with neuronal cells define the anatomy of the the pharynx (**Fig. 2d** and **Supplementary Fig. 2d**).

Another significant advance of STASCAN compared to the existing methods is that STASCAN enables accurate cell-type prediction solely based on corresponding spot images. We compared the performance of STASCAN, Tangram, and MUSE in predicting cell types when morphological images are provided while gene expression information is masked (**Fig. 2f**). STASCAN achieved precise cell annotation predictions which were consistent with those made when both image and gene expression data were avaiable. However, Tangram failed to predict cell types without gene expression data. Although MUSE achieved the characterization of cell clusters solely based on images, it was also disturbed by the absence of gene expression data, leading to incorrect predictions. For instance, MUSE identified two distinct cell clusters in the gut region that were disharmony with the manual annotations, and also failed to identify the pattern of neuronal cells in the pharynx region (**Fig. 2f**). This comparison highlights the superiority of STASCAN and provides the basis for its utilization in three designed application modules in the subsequent steps.

### STASCAN achieves super-resolution cellular patterns and improves 3D reconstruction in planarian

Next, we assessed the capabilities of STASCAN in different application modules using the planarian dataset^21^. When using Seurat and Cell2location to predict the spot cell types for pre-labeling, approximately half of the raw spots cannot be assigned with reliable cell labels (**Supplementary Fig. 2a** and **Methods**). This issue may be due to the noises generated by the complex signatures of gene expression in each spot, indicating the drawback of deconvolution in determining cell types (**Fig. 3a**). Choosing the other half of raw spots with credible labels as the prior spots to train the model, STASCAN achieved the reliable ability of cell type annotation based on images and depicted super-resolved cell distribution map (**Fig. 3a**). Firstly, STASCAN performed cell annotation for the unseen spots and demonstrated an enhanced resolution of cell distribution through combining both unseen spots and raw spots. The enhanced cell distribution map was highly consistent with H&E staining images, highlighting the relevant structures that were not shown at the raw resolution, such as the ventral nerve cord, genital chamber, pharynx, and contour (**Fig. 3a** and **Supplementary Fig. 2d**). Besides, it was highly consistent with the distribution of corresponding cell markers reported in previous literatures^23^ (**Fig. 3b** and **Supplementary Fig. 3a**).

**Fig. 3.**
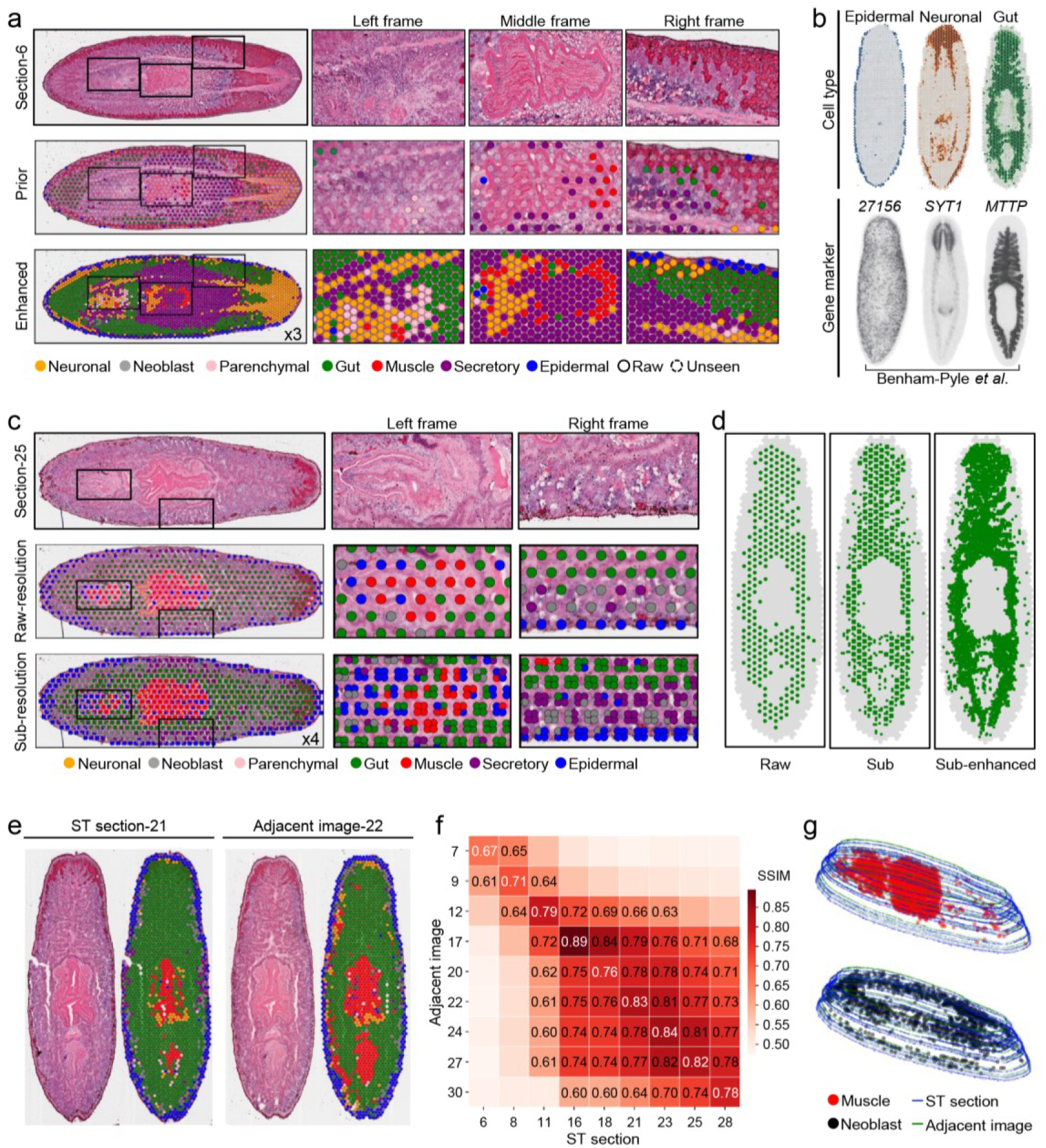
STASCAN provides comprehensive and multidimensional cell annotation in 10x Visium planarian dataset. **(a)** Tissue regions of the planarian in section 6. Left top, H&E staining image showing general morphology. Left center, spatial plot (x and y axis) covering the H&E staining image depicting distribution for the major cell types on the prior spots. Prior spots, the spots that are predicted to be the same cell types by both the Cell2location and Seurat at the raw resolution. Left bottom, spatial plot (x and y axis) covering the H&E staining image depicting distribution for the major cell types on the fully-tiled spots. The number in the bottom right indicates a 3 times improvement in cellual resolution over the raw resolution. Fully-tiled spots, containing both the raw spots and unseen spots. Raw spots, containing prior spots and other spots at the raw resolution. The latter were assigned with cell types by STASCAN. Unseen spots, the artificial spots among unmeasured areas, inferred with cell types by STASCAN. Right, enlarged view of the black framed area, arranged by left, middle, and right frame. **(b)** Visualizations of cell distribution for three major cell types and expression patterns of corresponding marker genes. Top, cell distribution annotated by STASCAN. Bottom, whole-mount in situ hybridization of classic gene markers (from published research^22^). **(c)** Tissue regions of the planarian in section 25. Left top, H&E staining image showing general morphology. Left center, spatial plot (x and y axis) covering H&E staining image depicting cell distribution for the major types annotated by STASCAN at raw resolution. Left bottom, spatial plot (x and y axis) covering the H&E staining image to decipher cell distribution for the major types annotated by STASCAN at sub-resolution. The number in the bottom right indicates a 4 times improvement in cellual resolution over the raw resolution. Right, enlarged view of the black framed area, arranged by left and right frame. **(d)** Spatial plots (*x* and *y* axis) depicting cell distribution for gut cells. Left, cell distribution predicted by STASCAN at raw resolution. Middle, cell distribution predicted by STASCAN at sub-resolution. Right, cell distribution predicted by STASCAN at sub-enhanced resolution. **(e)** Tissue region of planarian in section 21 and adjacent image 22. Left, H&E staining image and spatial plots (*x* and *y* axis) depicting cell distribution predicted by STASCAN of section 21. Right, H&E staining image and spatial plots (*x* and *y* axis) to decipher cell distribution predicted by STASCAN of adjacent image 22. **(f)** Heatmap showing the values of SSIM between the cell distribution of adjacent images and nether ST sections. Numbers on the coordinate axis represent the order of ST sections and adjacent images from the same consecutive sections set. **(g)** 3D spatial plots (x, y, and z axis) depicting 3D distribution of two selected cell types. Top, muscle cell. Bottom, neoblast cell. Both 3D models are reconstructed with aligned outlines and spots from the ST section and adjacent images.

Furthermore, STASCAN pinpointed the composition of cell mixtures and their distinct locations at sub-resolution, effectively distinguishing cell types of each subdivided spot, and displaying a more detailed distribution of fine-grained cells (**Fig. 3c**). For example, STASCAN sensitively allocated secretory and neoblast cells around the contour into sub-divided positions according to the morphological differences. STASCAN also identified muscle cells located at the junction of the pharynx and intestine at sub-resolution and were consistent with the biological priori information (**Fig. 3c**), which were not discovered at raw resolution from a group of gut cells. In addition, we utilized STASCAN to predict the enhanced sub-resolved distribution of gut cells, obtaining the fine-grained distribution of gut cells and reproducing the classical branching structure of the planarian intestinal tract (**Fig. 3d**). These results indicate that STASCAN significantly enhances cell granularity at sub-resolution, facilitating depiction of instrumental sub-structure with fine-grained cells.

Last but not least, STASCAN achieved the prediction of cell distribution in the unseen sections only by H&E images using the learn features of the adjacent ST sections (**Fig. 3e** and **Supplementary Fig. 3b**). In line with the biological interpretation that the cell distribution between two consecutive sections should be similar, the structure similarity index measure (SSIM) (ranging from 0.67 to 0.89) was linearly correlated with the spacing distance between adjacent images and ST sections, demonstrating the prediction accuracy of cell annotation for unseen sections (**Fig. 3f**, **Supplementary Fig. 3b** and **Methods**). Furthermore, we generated raw and unseen spots from ST sections and adjacent images, applied STASCAN to predict cell types for those spots, and then reconstructed 3D models for different structures with cellular patterns (**Fig. 3g**). The model displayed the cell distribution in three-dimensional, with improved cellular resolution in spatial and promoted utilization of staining images without ST sequencing.

### STASCAN identifies well-defined boundaries of distinct cell layers in the human intestinal tissue

To further evaluate the performance of STASCAN for ST datasets on different tissue architectures, we applied STASCAN to human intestinal datasets generated by 10× Visium technology. These datasets consisted of eight slides sampled at diverse sampling locations and time points^24^. We trained each slide using STASCAN with different sizes of prior spots, ranging from 297 to 1,551, and observed stable performances across all sizes (**Supplementary Fig. 4a-c**, **Supplementary Table 1** and **Methods**).

Next, we applied STASCAN to predict cell types of unseen spots. Compared with the cell distribution of prior spots annotated by other methods, STASCAN stratified cell populations to finer regional layers. For example, in comparison with other methods only roughly distinguishing the distribution of different cells, STASCAN was able to delineate the borders of cell layers among the intestinal epithelium, fibroblasts, and muscularis, greatly enhancing the cellular spatial patterns (**Fig. 4a** and **Supplementary Fig. 5**).

**Fig. 4.**
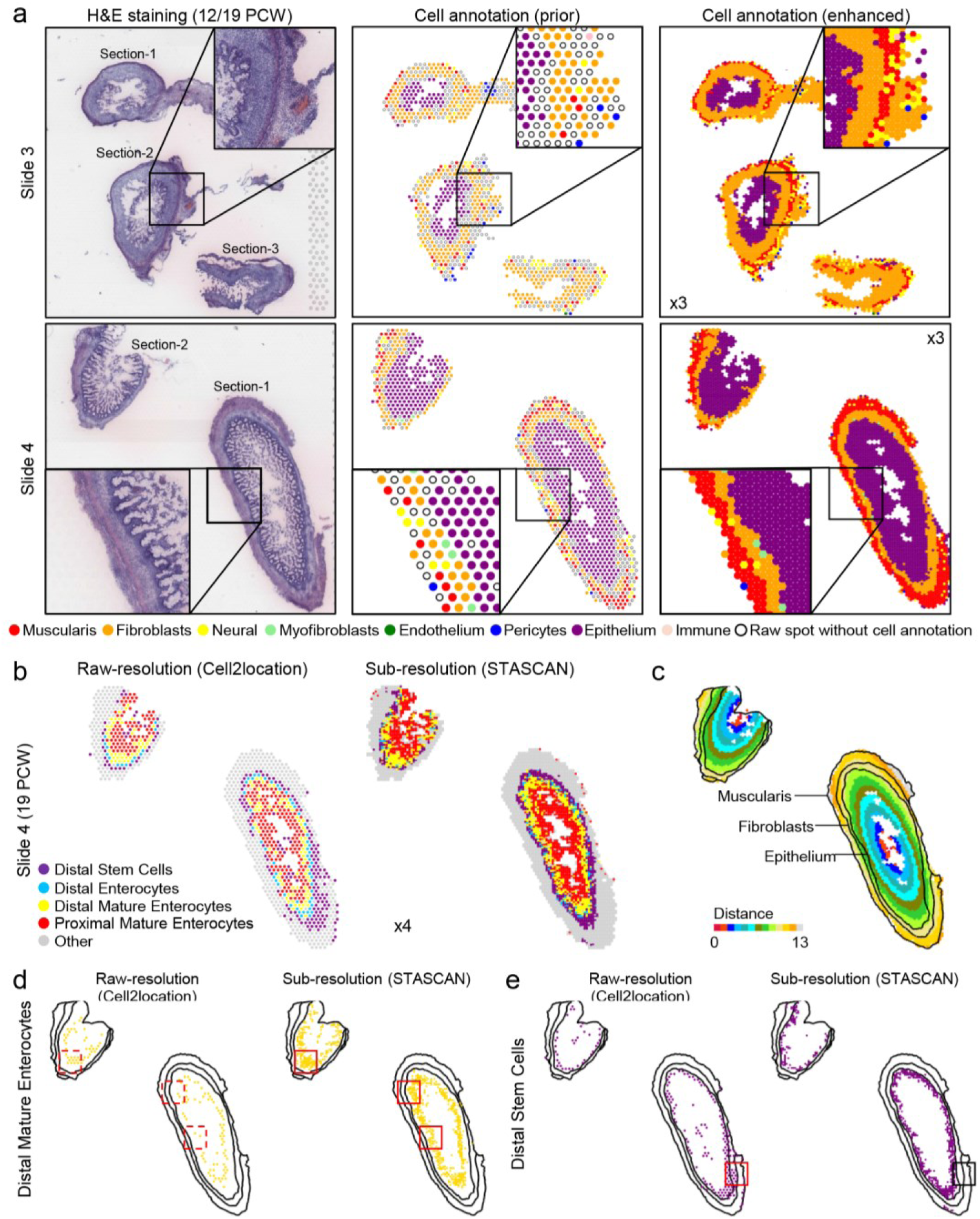
STASCAN depicts spatial layers of distinct cell types in 10x Visium human intestinal dataset. **(a)** Tissue regions of the human intestinal tissue for both the 12 PCW colon (slide 3) and 19 PCW colon (slide 4) respectively. Left, H&E staining image showing general morphology. Middle, spatial plot (x and y axis) depicting distribution for the major cell types of the prior spots. Right, spatial plot (x and y axis) depicting super-resolution cell distribution for the major cell types of the fully-tiled spots. Slide numbers are shown in columns. The numbers indicate a 3 times improvement in cellual resolution over the raw resolution. **(b)** Tissue regions of the human intestinal tissue for 19 PCW colon. Left, spatial plot (x and y axis) depicting cell distribution for epithelium subtypes inferred by Cell2location at raw resolution. Right, spatial plot (x and y axis) depicting cell distribution for epithelium subtypes inferred by STASCAN at sub-resolution. The number in the bottom left indicates a 4 times improvement in cellual resolution over the raw resolution. **(c)** Schematic of muscularis, fibroblasts, and epithelium layers, colored in gradients according to the distance from the spot to the tissue edge. **(d)** Spatial plots (*x* and *y* axis) depicting cell distribution for distal mature enterocytes. Left, cell distribution predicted by Cell2location at raw resolution. Right, cell distribution inferred by STASCAN at sub-resolution. The black lines represent the schematic of different morphological layers. **(e)** Spatial plots (*x* and *y* axis) depicting cell distribution for distal stem cells. Left, cell distribution inferred by Cell2location at raw resolution. Right, cell distribution inferred by STASCAN at sub-resolution. The black lines represent the schematic of different morphological layers.

Then, we used STASCAN to draw the spatial distribution map of fine cell subtypes in the human intestinal tissue (**Fig. 4b**). Actually, we labeled three anatomical layers of the intestinal tissue related to the morphological structures of H&E staining, including muscularis, fibroblasts and epithelium layers, listed based on their distance to the intestinal edge (**Fig. 4c**). When evaluating four epithelium subtypes occupying an absolute proportion of the epithelium layer, we found that compared to the results of alternative method at raw-resolution, STASCAN not only highlights the precise distribution of these subtypes but also accurately locates the positions of distinct subtype cells (**Fig. 4b, d, e** and **Supplementary Fig. 6a**). For instance, distal epithelium subtype cells tend to gather closer to the boundary of the epithelium layer and the fibroblast layer, and proximal epithelium subtype cells are prone to assemble to the surface of the epithelium layer at sub-resolution. Besides, distal stem cells were correctly predicted to be located in the epithelium layer at sub-resolution, however, at raw-resolution a part of distal stem cells were abnormally predicted to be located in the fibroblasts layer (**Fig. 4d, e** and **Supplementary Fig. 6b**).

In addition, we performed STASCAN on a pair of identical and adjacent sections of the intestinal tissue to valid the cell annotation for unseen sections (**Methods**), and the high correlation between them further confirmed the accuracy and reliability of the prediction (**Supplementary Fig. 7a, b**).

### STASCAN uncovers a novel structure in the human lung tissue

Despite the limitations of raw spatial resolution of ST technologies, STASCAN provides assistance in enhancing cellular pattern and rediscovering the micrometer-scale structure. Here, we applied STASCAN on the 10× Visium human lung dataset^25^, which sampled from the proximal airway. We previously redefined 13 reference cell types to bette illustrate the organizational structure and annotated 822 ST spots with seven dominant cell types to train the STASCAN (**Supplementary Fig. 8a-e**, **Supplementary Table 1** and **Methods**). With enhanced resolution, STASCAN showed more precise cellular and structural patterns of human lung tissue. Besides, we observed that STASCAN sensitively identified a micrometer-scale oval-shaped structure that was highly consistent with the H&E staining images, which was confirmed as the smooth muscle bundles adjacent to the tracheal wall. However, this structure was not evident at the raw resolution of prior spots, highlighting the capability of STASCAN to reveal refined structures of spatial regions (**Fig. 5a** and **Supplementary Fig. 8c**).

**Fig. 5.**
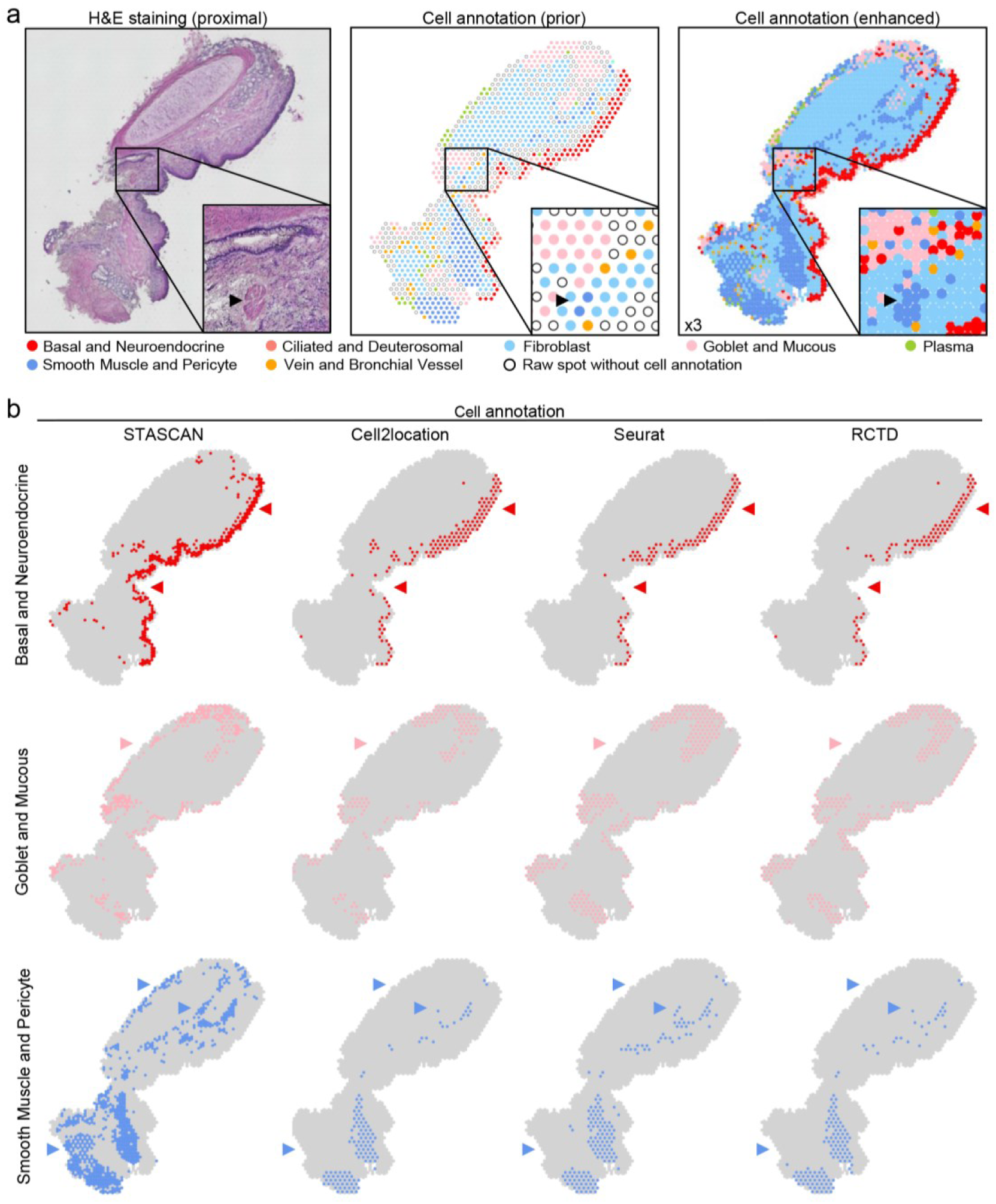
STASCAN demonstrates the special structure from 10x Visium human lung data. **(a)** Tissue regions of the human proximal lung. Left, H&E staining image showing general morphology. Middle, spatial plot (*x* and *y* axis) depicting cell distribution for the major types on prior spots. Right, spatial plot (x and y axis) depicting super-resolution cell distribution for the major types on the fully-tiled spots. The number in the bottom left indicates a 3 times improvement in cellual resolution over the raw resolution. **(b)** Spatial plots (*x* and *y* axis) depicting cell distribution for the three major types. Cell types are shown in columns, and alternative methods (STASCAN, Cell2locaiton, Seurat, and RCTD) used to annotate the corresponding cell type are shown in rows.

Furthermore, we compared the ability of STASCAN with other methods in revealing tissue structures with cellular patterns. The results showed that STASCAN could clearly depict the silhouette of the airway with basal and neuroendocrine cells and the cricoid cartilage structure surrounded by goblet cells, mucous cells, smooth muscle cells, pericytes, etc. Moreover, the smooth muscle tissue traced by smooth muscle cells and pericytes in the left bottom of the slide was only identified by STASCAN (**Fig. 5b** and **Supplementary Fig. 8c**). Briefly, compared with the cell distribution pattern identified by other methods, STASCAN displayed superior advantages in identifying and characterizing spatial specific structures, which better reflects the anatomical structure after imputation.

### STASCAN depicts the pathological spatial structural variations of human cardiac tissue after myocardial infarction

To explore whether further improved functional applications in ST data analysis can be achieved by STASCAN, we adopted this approach to reanalyze the 10× Visium human cardiac datasets^26^, which included 17 slides from normal hearts and the pathological ones after myocardial infarction (**Supplementary Figs. 9a,b, 10**, **Supplementary Table 1** and **Methods**). We first grouped these slides according to their sampling regions^26^, including normal non-transplanted donor hearts as controls, necrotic (ischaemic zone and border zone), and unaffected regions (remote zone), and regions at later stages after myocardial infarction (fibrotic zone). After performing STASCAN to enhance the spatial cell distribution, we focused on two slides with a lot of missing spots that were filtered out due to scanty genes and unique molecular identifiers (UMI) measured in the original literature^26^ (**Fig.6a** and **Supplementary Figs. 11-15a**). In these two slides, STASCAN not only more accurately depicted the cell distribution pattern of tissue structures, but also predicted the potential cell distributions in the missing areas only from images. Especially for the serious missing in the slide_ACH0010 sampled from the ischaemic zone, STASCAN better imputed the reasonable diffusion of the cardiomyocyte, fibroblast, and myeloid cells in line with the histological morphology (**Fig. 6a**). The proximity of the latter two cells indicated a strong dependence between them on the areas of immune cell infiltration and scar formation^26^.

**Fig. 6.**
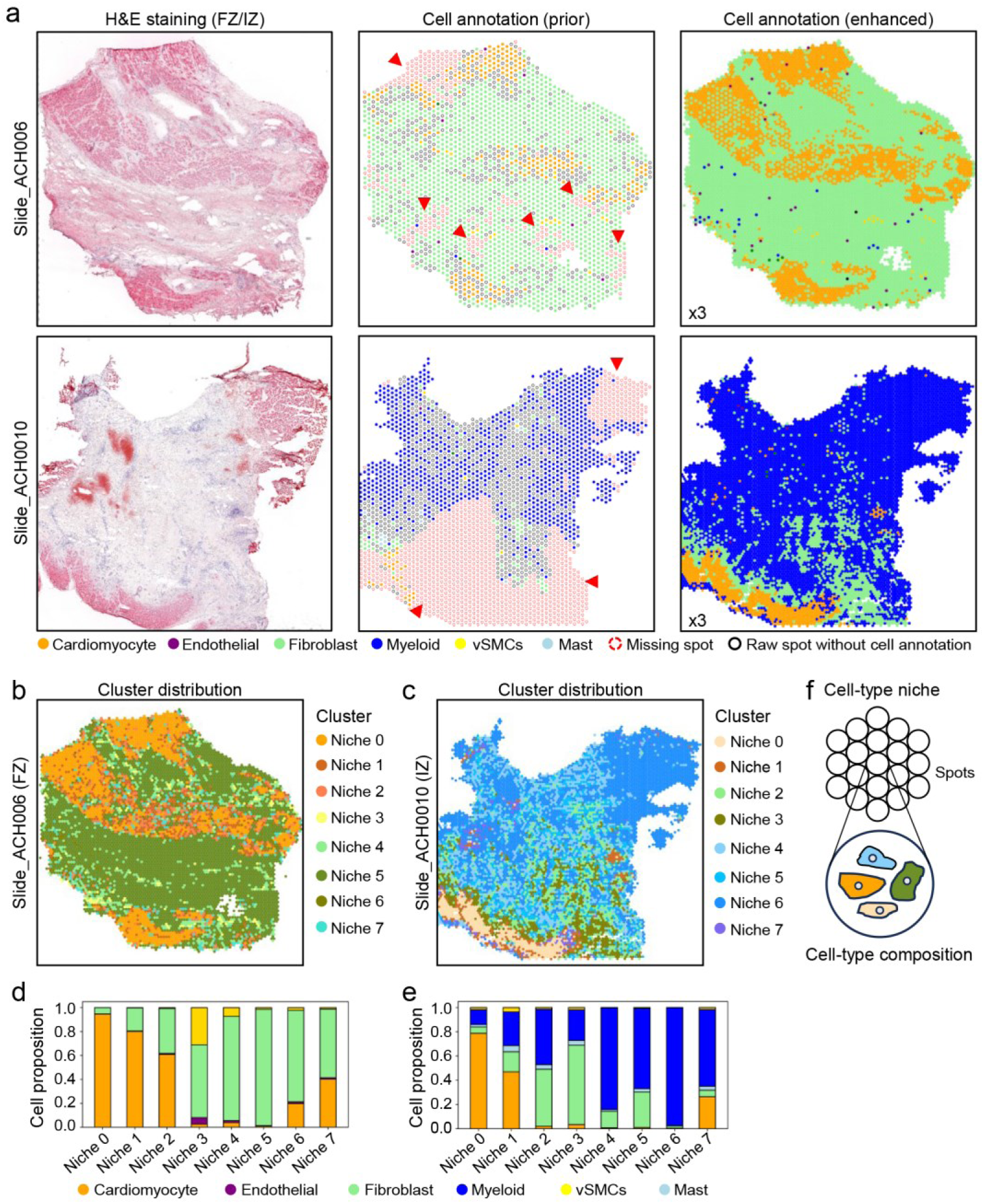
STASCAN reveals the cell-type niches in 10x Visium human cardiac data. **(a)** Tissue regions of the human cardiac tissue from the fibrotic zone (Slide_ACH006) and ischaemic zone (Slide_ACH0010) after myocardial infarction. Left, H&E staining image showing general morphology. Middle, spatial plot (*x* and *y* axis) depicting cell distribution for the major cell types on prior spots. Right, spatial plot (x and y axis) depicting super-resolution cell distribution for the major cell types on the fully-tiled spots. The numbers in the bottom left indicate a 3 times improvement in cellual resolution over the raw resolution. Slide numbers are shown in columns. Missing spots, spots removed from raw spots in the original literature due to the small number of measured genes and UMIs. FZ, fibrotic zone. IZ, ischaemic zone. **(b)** Spatial plots (*x* and *y* axis) depicting spatial distribution for cell-type niches in slide_ACH006, which are assigned by unsupervised clustering of spots with the cell-type compositions. **(c)** Spatial plots (*x* and *y* axis) depicting spatial distribution for cell-type niches in slide_ACH0010, which are assigned by unsupervised clustering of spots with the cell-type compositions. **(d)** Bar plot showing the cell proposition of cell-type niches in slide_ACH006. **(e)** Bar plot showing the cell proposition of cell-type niches in slide_ACH0010. **(f)** Schematic of cell-type niches.

We then explore the spatial structural variations in these two slides by performing unsupervised clustering for spots based on the composition of cell annotations predicted by STASCAN, and then mapped the clusters, defined as cell-type niches, to the spatial regions (**Fig. 6b, c, f** and **Supplementary Figs. 11-15b**). Through redrawing the spatial distributions for these cell-type niches, the cardiac tissue manifested more delicate spatial patterns compared with the dominant cell annotation, which were consistent with histological morphology and detailed structural variations observed during physiological and pathological processes.

Besides, these cell-type niches based on diverse cell propositions revealed more elaborate cell interacting microenvironments with potential biological insights (**Fig. 6d, e** and **Supplementary Figs. 11-15c**). For example, we observed myogenic cell-type niches (0, 1, and 2) mainly displaying characteristics of cardiomyocyte cells, and fibrotic cell-type niches (3, 4, 5, 6, and 7) mainly presenting characteristics of fibroblast cells, in the slide_ACH006 sampled from the fibrotic zones. On the aspect of spatial distributions, the myogenic cell-type niches could jointly characterize the myocardial structure, and the fibrotic cell-type niches distinguished by the proprotion of fibroblast cells indicated different fibrosis processes during the lesion. Especially, there was a measure of characteristics of vSMCs and endothelial cells in niches 3 and 4 for depicting the spatial structure of the cardiac vasculature. In addition, in slide_ACH0010, we observed inflammatory cell-type niches (4, 5, 6, and 7) which mainly exhibit characteristics of myeloid and mast cells, apart from myogenic cell-type niches (0 and 1) and fibrotic cell-type niches (2 and 3). These three types of niches took up distinct spatial regions, but there was niche 7 located at the intersection, in line with the proposition of cardiomyocyte, fibroblast, myeloid, and mast cells in the niche 7. Especially, niches 2, 3, 4, and 5 showed co-enrichment between myeloid and fibroblast cells, in accordance with the role of macrophages in fibroblast activation^27^ and fibroblast cells in macrophage attraction^28^. Overall, STASCAN expands the application of niche distribution and provides better insights into understanding cellular microenvironment interactions.

### STASCAN deciphers the intricate tissue organization throughout the developmental stages of the mouse brain

We further test whether STASCAN is also applicable in ST data derived from various technologies. We first employed STASCAN on an embryonic mouse brain dataset from MISAR-seq^29^, a microfluidic indexing-based spatial technology motivated by DBiT-seq^7^ with both high-quality image and sequencing data (**Supplementary Fig. 16**, **Supplementary Table 1** and **Methods**). Notably, although the H&E images adopted in this dataset were obtained from the adjacent tissue slide which causes a partial disharmony between the actual gene expression pattern and morphological images, STASCAN still achieved excellent results. When compared with the cellular distribution annotated by RCTD^29^, STASCAN significantly improved the cellular resolution with highlighted characteristics of tissue structures (**Fig. 7a** and **Supplementary Fig. 16**). For example, the enhanced distribution pattern of forebrain GABAergic neurons was associated with the subpallium, and a group of the forebrain glutamatergic and cortical or hippocampal glutamatergic neurons spotlighted the dorsal pallium of the forebrain at enhanced resolution (**Fig. 7a**).

**Fig. 7.**
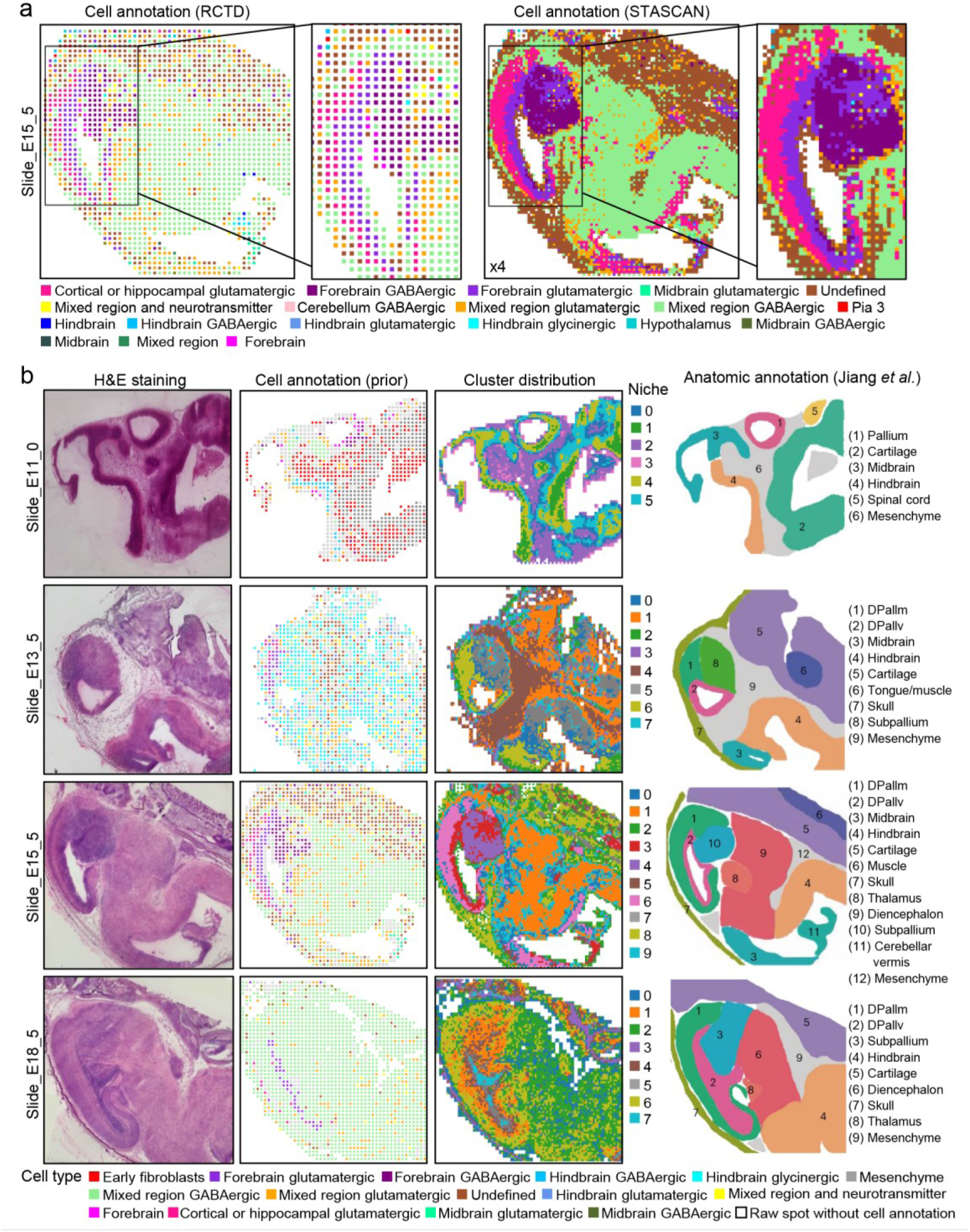
STASCAN revealed major anatomical tissue regions in mouse brain dataset generated from microfluidic technologies. **(a)** Tissue regions of the E15.5 embryonic mouse brain tissue. Left, spatial plot (*x* and *y* axis) depicting cell distribution annotated by RCTD. Right, spatial plot (x and y axis) depicting super-resolution cell distribution annotated by STASCAN. The number in the bottom left indicates a 4 times improvement in cellual resolution over the raw resolution. **(b)** Tissue regions of the embryonic mouse brain tissues across development stages. Left, H&E staining image showing general morphology. Middle left, spatial plot (x and y axis) depicting cell distribution for the major cell types on prior spots. Middle right, spatial plot (x and y axis) depicting spatial distribution for cell-type niches, which are assigned by unsupervised clustering of spots with the cell-type compositions. Slide numbers are shown in columns. Right, manual anatomic annotation of major tissue regions for different mouse brain stages from original literature^29^. DPallm, mantle zone of the dorsal pallium. DPallv, ventricular zone of the dorsal pallium.

Furthermore, we respectively generated cell-type niches for different development stages of mouse brain tissue and mapped them to spatial regions (**Fig. 7b** and **Methods**). Using the manual anatomical annotations of major tissue organizations from H&E images as the ground truth^29^ (**Fig. 7b**), we compared the cluster distribution of cell-type niches to the raw-resolution distributions of cell annotations on prior spots. The cluster distribution of cell-type niches generated by STASCAN more remarkably recapitulates the tissue organization in the developing mouse brain than the ones from raw-resolution. Especially for the E18.5 embryonic mouse brain tissue, in contrast to cell annotation at raw resolution which showed an unrecognized organization in nearly entire brain region, STASCAN clearly defined the major tissue domains by cell-type niches clustering (**Fig. 7b**). Collectively, these results illustrate the strength of STASCAN in highlighting tissue structures and redrawing finer organizations using ST data from various technologies.

## Discussion

Although current spatial transcriptomics has achieved remarkable progress in deciphering the distribution and interaction of cells in tissues, the spatial resolution limitations hinder its broader application. The development of computational algorithms is crucial for the analysis of spatial transcriptomic data. Here, we have developed STASCAN, a tool that integrates histological images and spatial gene expression to depict comprehensive cell atlases with enhanced spatial resolution.

Compared with the traditional image-based CNN models, STASCAN integrates gene expression and image information, in which gene expression helps to automatically label spots used for training without manual annotation. Meanwhile, images extend a new perspective and supplement the information that sequencing cannot provide. Additionally, STASCAN can accurately extract features and improve the accuracy of the model through transfer learning and pseudo-labeling. Thus, STASCAN effectively resolves images and uses the image information as the main reference for cell type determination. This approach helps to predict cell distribution and increases the number of spots in the point clould representing cells, ultimately facilitating the construction of fine-resolved 3D spatial cell distribution maps.

Through fully utilizing both sequencing and image data, STASCAN not only accurately annotates the cell types on raw ST spots but also enables the prediction of cell distribution in the uncharted areas across tissues. Besides, STASCAN fully utilizes the image information of continuous sections to construct 3D tissue models without expensive ST experimental costs, laying the foundations for expanding the future frontier in 3D cellular atlas. We performed STASCAN in different species and tissues and observed a substantial advance over current methods in depicting spatial cellular patterns and tissue structures. In planarian, STASCAN successfully predicted cell distribution of unseen spots and sections in the uncharted areas, leading to the construction of more detailed 3D cell distribution models. In addition, we found that STASCAN discovers precise tissue structures with enhanced cell annotation. In human lung tissue, STASCAN enhanced spatial resolution and identified a micrometer-scale structure that was made up of a group of smooth muscle cells located near the tracheal wall and involved in the regulation of intra-pulmonary airway caliber^30^, while the raw resolution are too coarse to resolve the same specific structure.

The low resolution of NGS-based spatial transcriptomics is also attributed to the spacing of captured spots, which are not small enough to achieve single-cell resolution^5^. Most current methods estimate the proportion or abundance of each cell type from the cell mixtures on the capture spots but are incapable of assigning each cell into exact locations within each spot. Although Tangram provides a computer vision module that segments cell nuclei based on histological images and predicts the cell type for each cell in the spot, it performs random cell assignment for the segmented maskings within the spot and still cannot definitely resolve cell position^17^. In comparison, STASCAN introduces a predicting module at sub-resolution, allowing the assignment of cell types to sub-divided spots based on the tissue morphological features corresponding to the exact positions. In human intestinal tissue, STASCAN precisely pinpointed the fine-grained cell subtypes between distinct cell layers. The advantages of STASCAN enable it to have a broader range of applications. For example, STASCAN can effectively predict cell distribution of some regions with missing sequencing data due to a low number of captured UMIs and genes. In human cardiac tissues, STASCAN could fill in the missing data, and help to display an extensive spatial proximity distribution of fibroblast and myeloid cells in the vicinity of scar tissue. This provides a better understanding of the relationship between fibroblasts and immune cells in myocardial infarction, such as immune factors in stimulating fibroblast transformation^31^. Additionally, the cell proposition annotations generated by STASCAN is supported to cluster different cell-type niches, providing novel insights into the microenvironment at the cellular resolution. In human cardiac tissues, STASCAN generated distinct niches including inflammatory, myogenic, and fibrotic cell-type niches, with the best representation of the dynamic diversity of cells in the microenvironment during myocardial infarction. In embryonic mouse brain tissue, STASCAN clearly revealed major anatomical tissue organizations across the development stages, with gradually increased complexity of brain structures during brain development.

Logically, STASCAN can be employed in all ST technologies that utilize microarrays, microfluidics, or similar designs, to annotate cell types for unseen spots. It can also be used in other NGS-based ST technologies to annotate cell types for subdivided spots and unseen sections, only if the technology provides both H&E staining images and transcriptional data. Due to the limitation in collecting high-quality H&E staining images of published ST datasets, we only applied STASCAN in ST datasets generated by the 10× Visium and MISAR-seq technologies, which respectively stands for microarray-based and microfluidic-based technologies. Nevertheless, the technical framework of STASCAN is well-designed and provides convenient interfaces for further implementation in other NGS-based ST technologies.

In summary, STASCAN is a universal and precise cell type prediction method, which integrates image and expression information to determine complex spatial distributions in tissues. The diverse applications of STASCAN have demonstrated its superior performance in enhancing spatial resolution and discovering novel structures, offering potential for resolving cell type distribution in various types of tissues across different stages of development, regeneration, and disease. The positional relationships of different cell types determined by STASCAN provide new insights into cell crosstalk critical in orchestrating organismal development and homeostasis. Furthermore, STASCAN fully utilizes more easily acquired image datasets under various biological conditions, and can potentially be used to infer subtypes of pathological cells and further link spatial cellular distribution to disease diagnosis cost-efficiently, through extensive data training, eliminating the need for sequencing.

