## Supplemental for "STASCAN deciphers fine-resolution cell-distribution maps in spatial transcriptomics by deep learning" (from another version of this preprint: version 1, 2023-09-04)

**Affiliations**

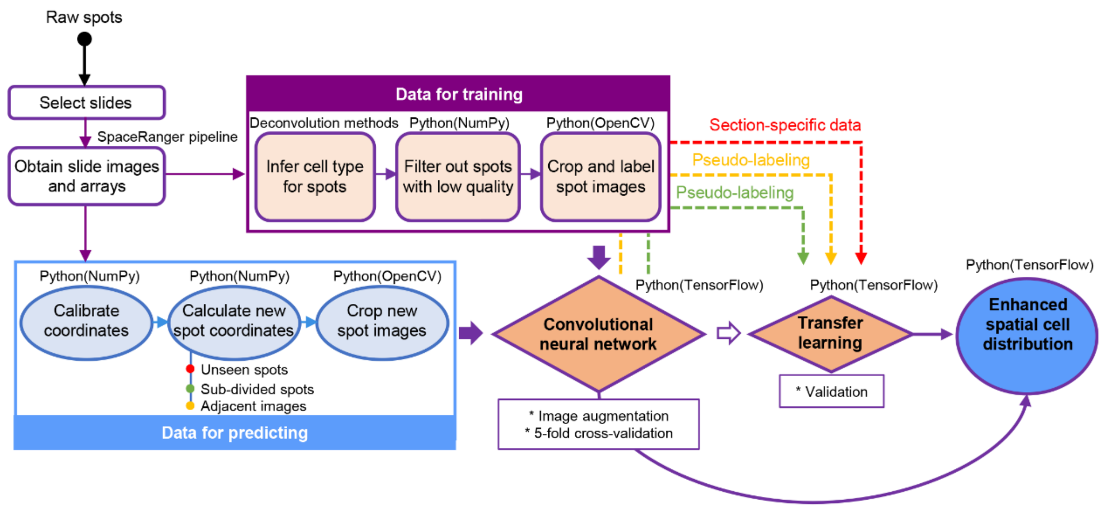


**Figure S1. Workflow of STASCAN.** The diagram consists of the process of data preparation, model training and prediction.

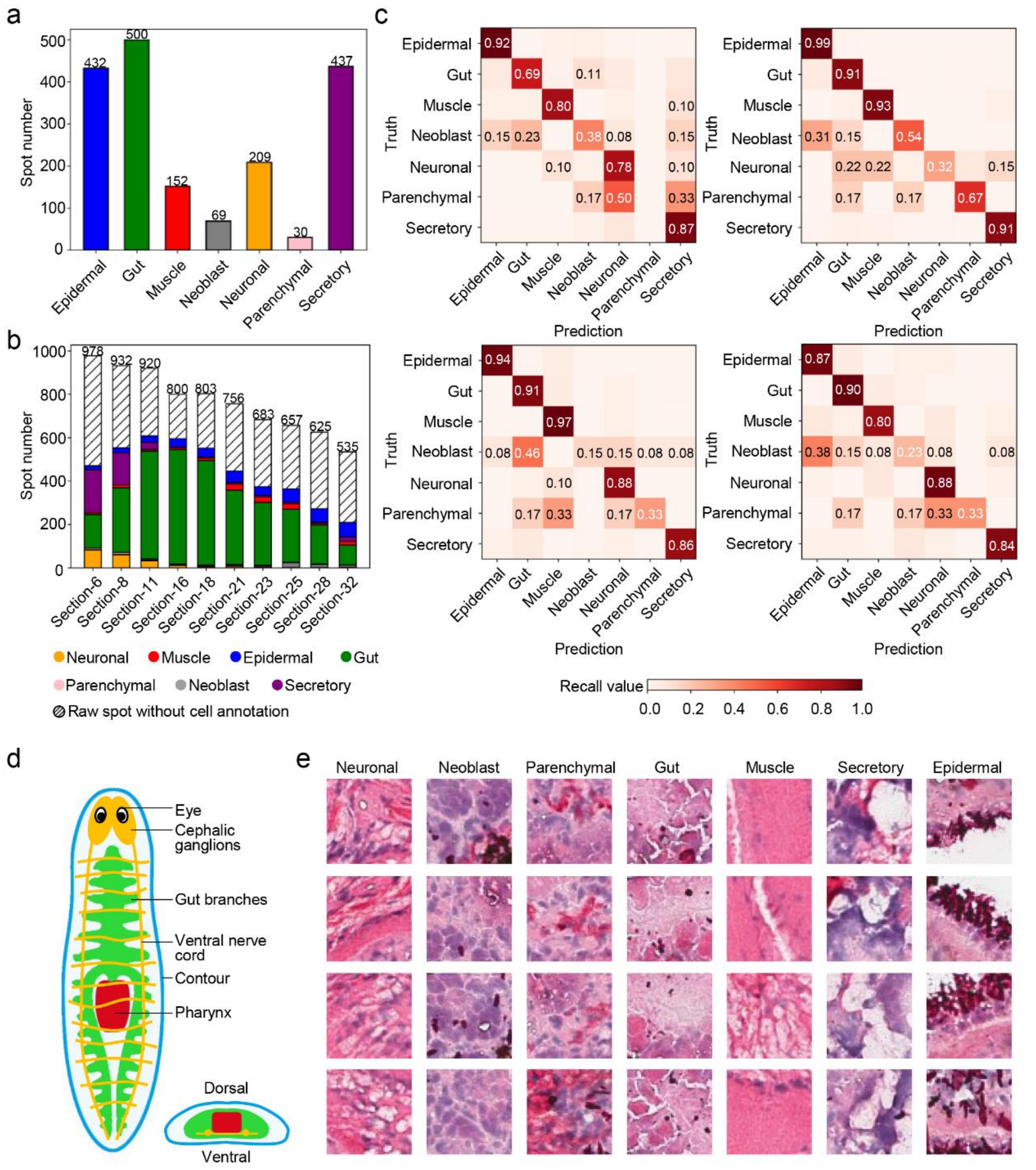

**Figure S2. Preparation and evaluation of the base model of STASCAN on the planarian dataset.**

**(a)** Bar plot showing the numbers of prior spots for each cell type on the base model in the planarian dataset.

**(b)** Bar plot showing the numbers of prior spots for the base model and the numbers of raw spots for each section in the planarian dataset. The bars filled with dashed lines represent the number of raw spots without cell annotation.

**(c)** Heatmaps showing the other confusion matrix of the prediction and truth of the base model in the planarian dataset with the five-fold cross-validation. The number in each box represents the recall value.

**(d)** Schematic representation showing the major tissue structure of the planarian.

**(e)** Examples of each major cell type with the corresponding image annotated by STASACN. The size of an image is equal to the diameter of a spot.

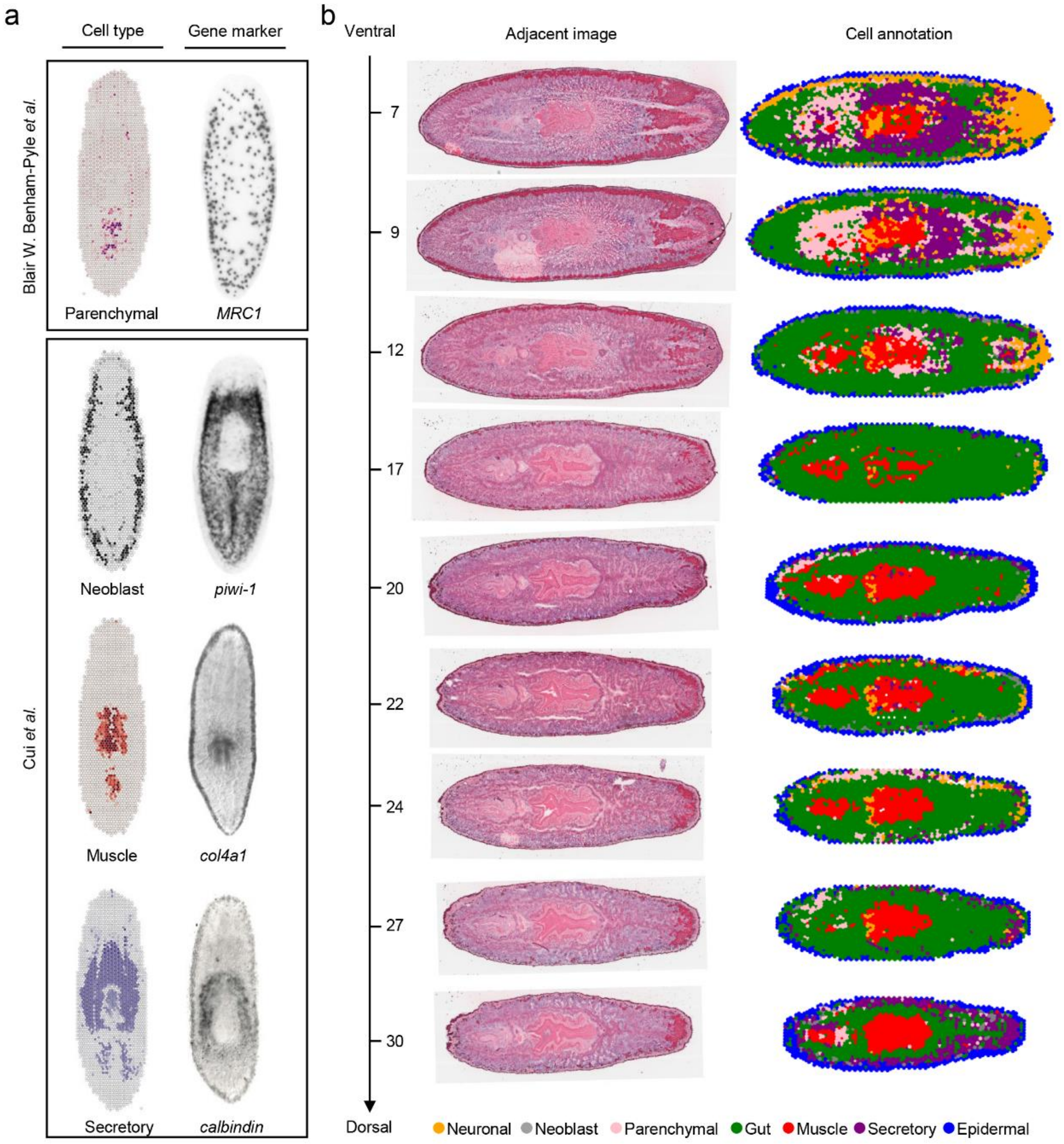


**Figure S3. Verification of STASCAN for unseen spots and unseen sections in the planarian dataset.**

**(a)** Visualization of cell distribution for the four major types and patterns of corresponding marker gene expression.

**(b)** Tissue regions of planarians in all adjacent images. Left, H&E staining of adjacent images. Right, spatial plots (x and y axis) depicting cell distribution predicted by STASCAN.

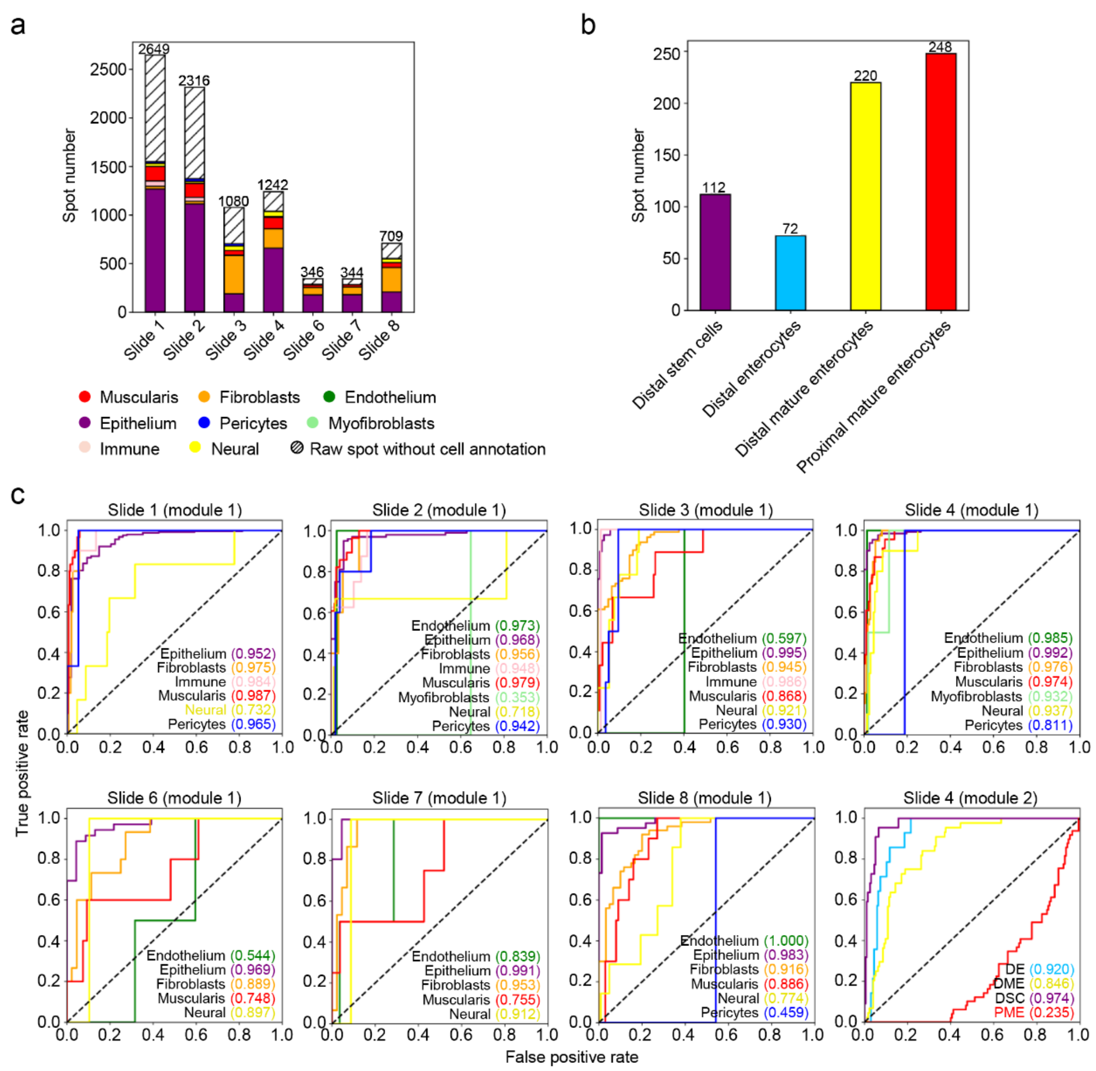


**Figure S4. Preparation and evaluation of STASCAN on the human intestinal dataset.**

**(a)** Bar plot showing the number of prior spots in each slide used for cell annotation models training for unseen spots. The bars filled with dashed lines represent the number of raw spots without cell annotation.

**(b)** Bar plot showing the number of prior spots on each cell type for the cell annotation model training for subdivided spots in the human intestinal dataset.

**(c)** The ROC curves of each cell type for models in the human intestinal dataset. The number in each pair of parentheses represents the AUC value. Module 1, trained for unseen spots. Module 2, trained for subdivided spots.

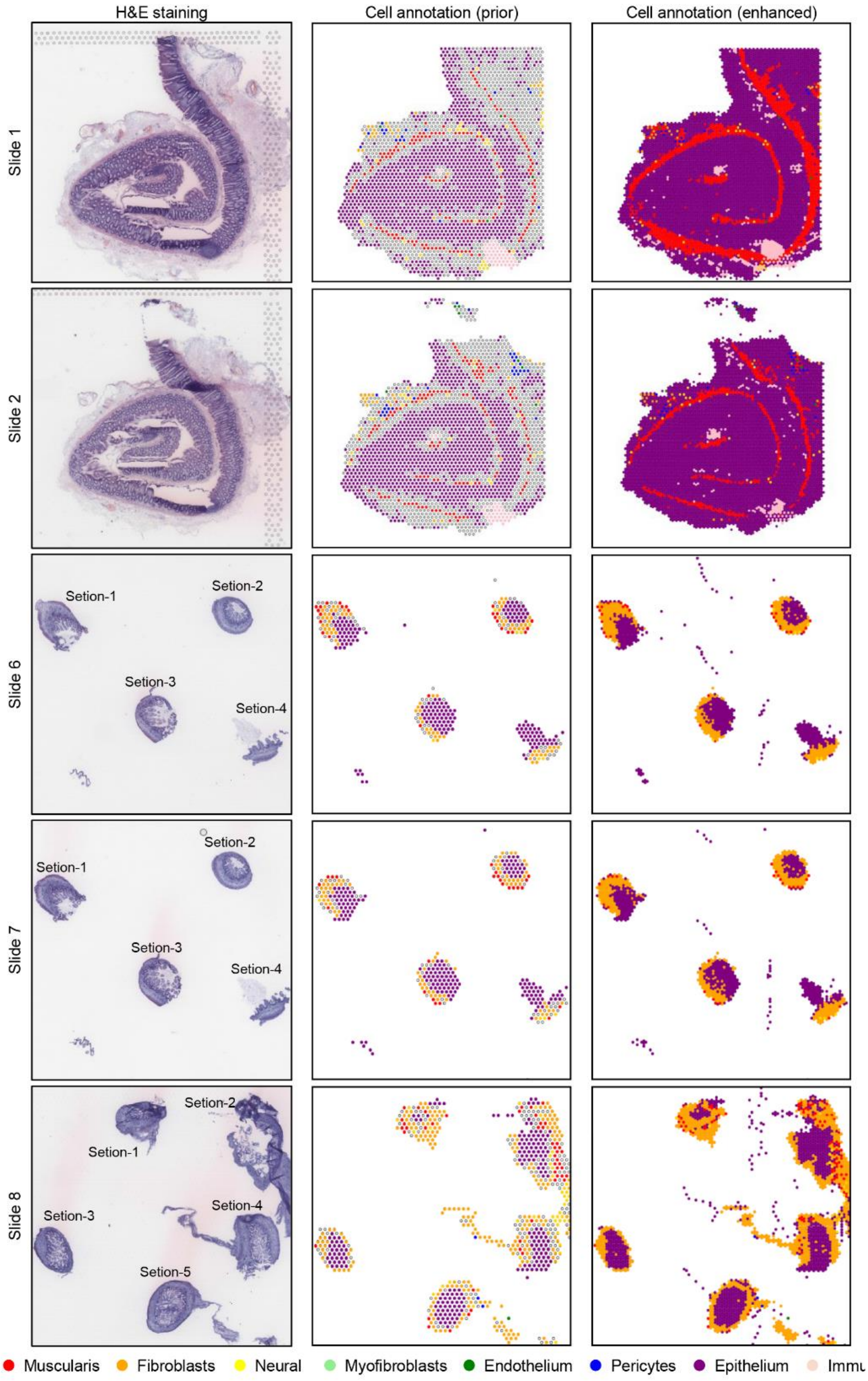


**Figure S5. Super-resolution cell distribution by STASCAN on the human intestinal dataset.** Tissue regions of other slides in the human intestine. Left, H&E staining image showing general morphology. Middle, spatial plot (x and y axis) depicting cell distribution for the major cell types on the prior spots. Right, spatial plot (x and y axis) depicting super-resolution cell distribution for the major cell types on the fully-tiled spots.

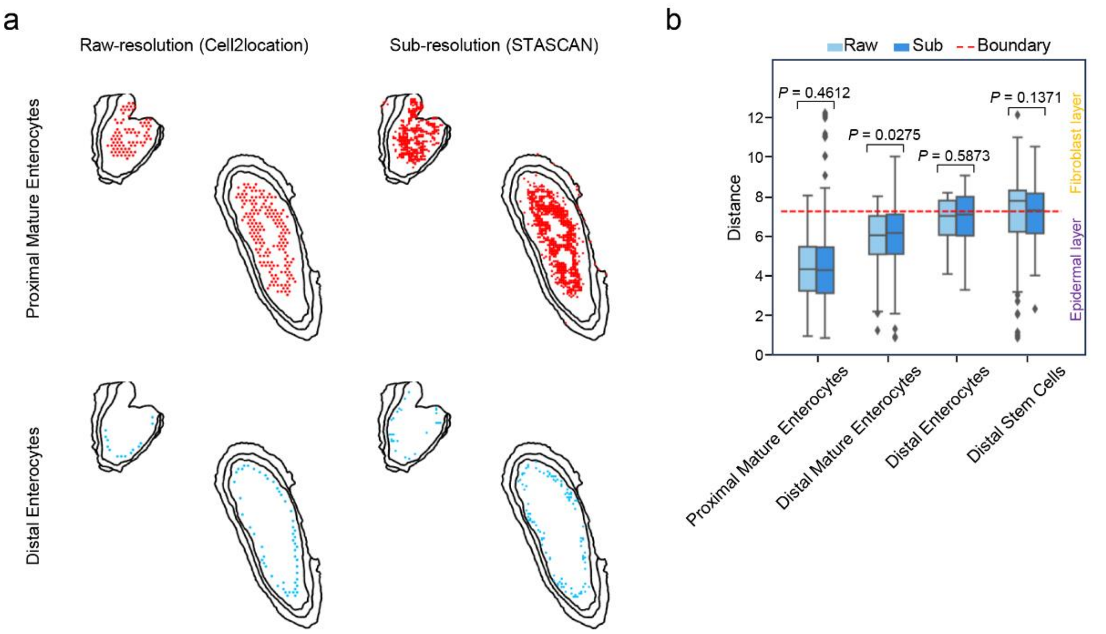


**Figure S6. Sub-resolution cell distribution of epithelium subtypes by STASCAN on the human intestinal dataset.**

**(a)** Spatial plots (x and y axis) depicting cell distribution for the other two epithelium subtypes. Left, cell distribution predicted by Cell2location at raw resolution. Right, cell distribution predicted by STASCAN at sub-resolution. Cell types are shown in columns. The black lines represent the schematic of different morphological layers.

**(b)** Box plot showing the distance score of epithelium subtypes at raw-resolution (n = 261, 121, 66, 189) and at sub-resolution (n = 982, 665, 126, 533), respectively. The red dashed line represents the average distance score of the boundary between the epidermal layer and the fibroblast layer. *P* value is calculated using the two-sided Wilcoxon signed-rank test.

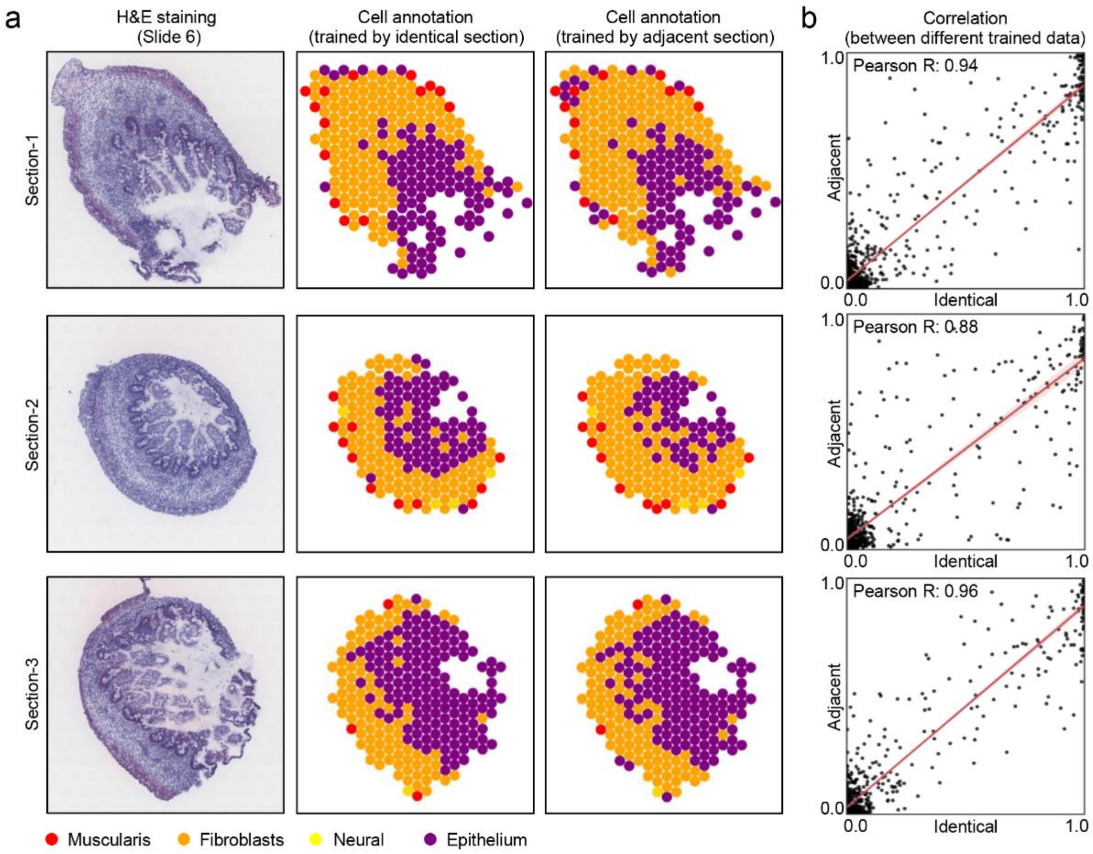


**Figure S7. Verification of STASCAN for unseen sections in the human intestinal dataset.**

**(a)** Tissue regions of slide 6 on the human intestinal dataset. Left, H&E staining image showing general morphology. Middle, spatial plot (x and y axis) depicting super-resolution cell distribution for the major cell types predicted from the model trained by the identical section spots. Right, spatial plot (x and y axis) depicting super-resolution cell distribution for the major cell types from the model trained by the adjacent section spots.

**(b)** Scatter plots showing the Pearson correlations between cell-type predictions obtained from the model trained by identical section spots and the model trained by adjacent section spots respectively.

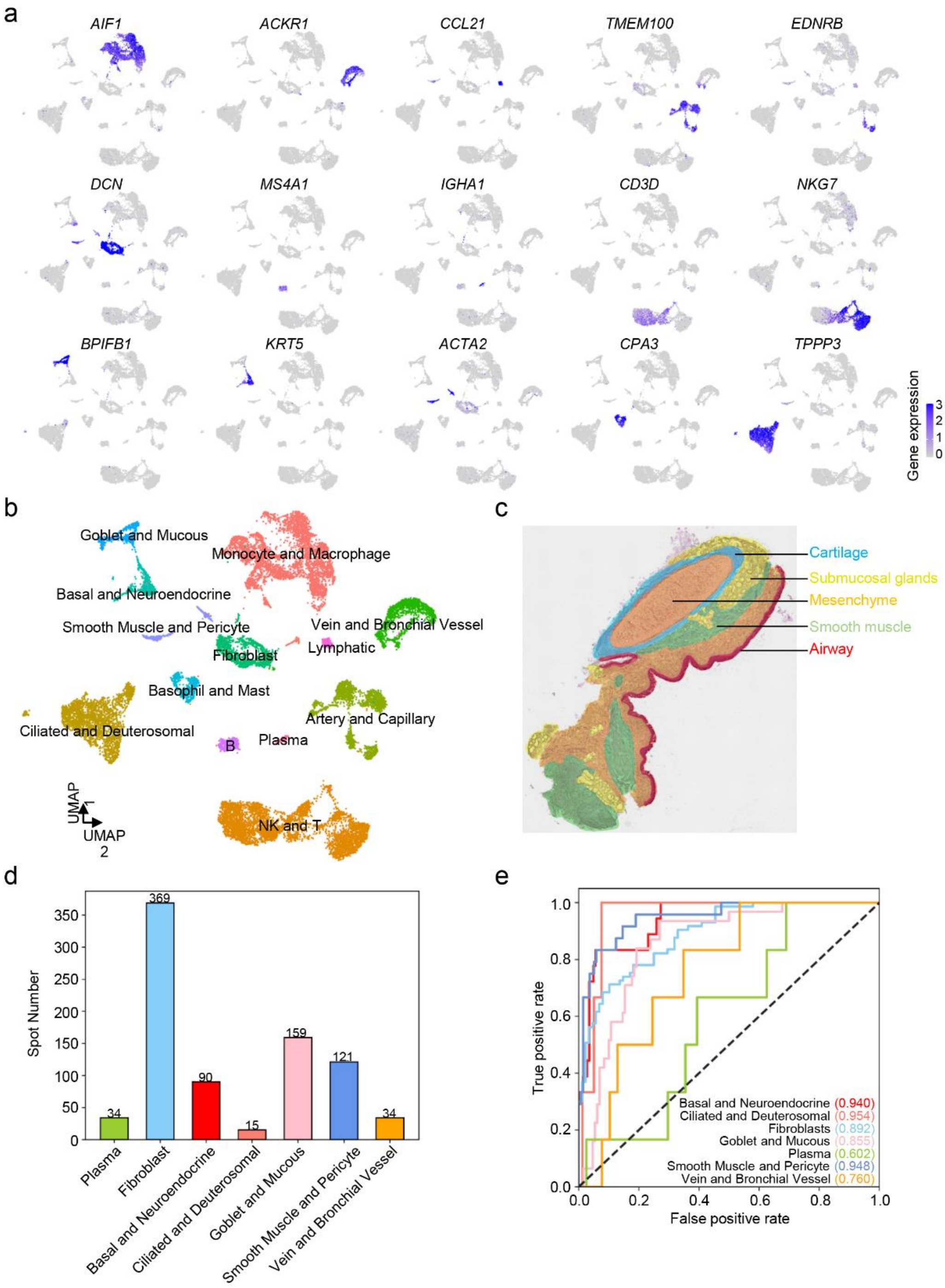


**Figure S8. Preparation and evaluation of STASCAN on the human lung dataset.**

**(a)** UMAP plot demonstrating the expression of marker genes for redefined clusters in the single-cell reference data.

**(b)** UMAP plot demonstrating the annotation of cell types for redefined clusters in the single-cell reference data.

**(c)** Schematic representation showing the major tissue structure of the human lung.

**(d)** Bar plot showing the number of prior spots used for training STASCAN for unseen spots in the human lung dataset, separately by cell types.

**(e)** The ROC curves of each cell type by STASCAN on the human lung dataset. The number in each pair of parentheses represent the AUC value.

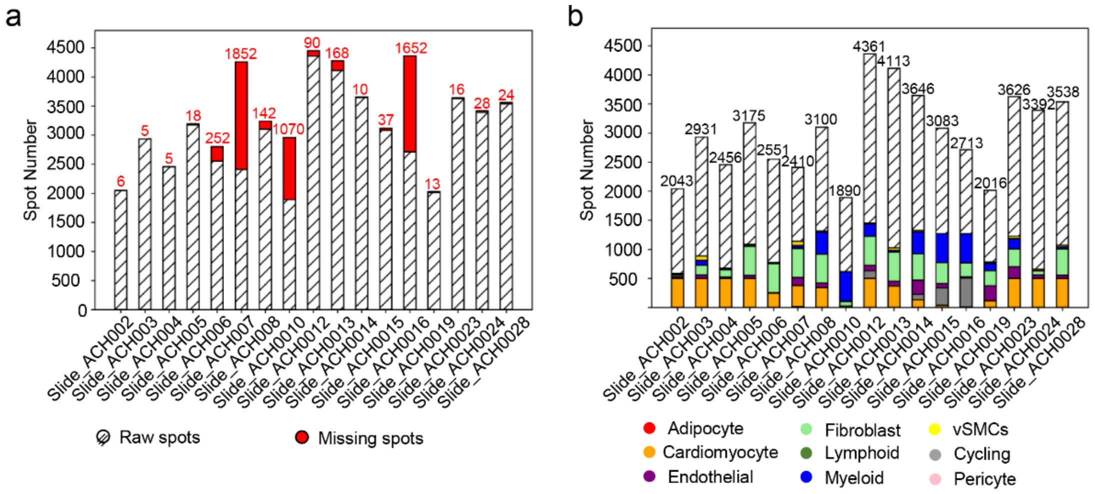


**Figure S9. Preparation of the human cardiac dataset.**

**(a)** Bar plot showing the number of raw spots and missing spots for each slide in the human cardiac dataset. The bars filled with dashed lines represent the number of raw spots on each slide. The red bars and numbers represent the number of missing spots.

**(b)** Bar plot showing the number of prior spots for training STASCAN for unseen spots and the number of raw spots for each slide in the human cardiac dataset. The bars filled with dashed lines represent the number of raw spots on each slide.

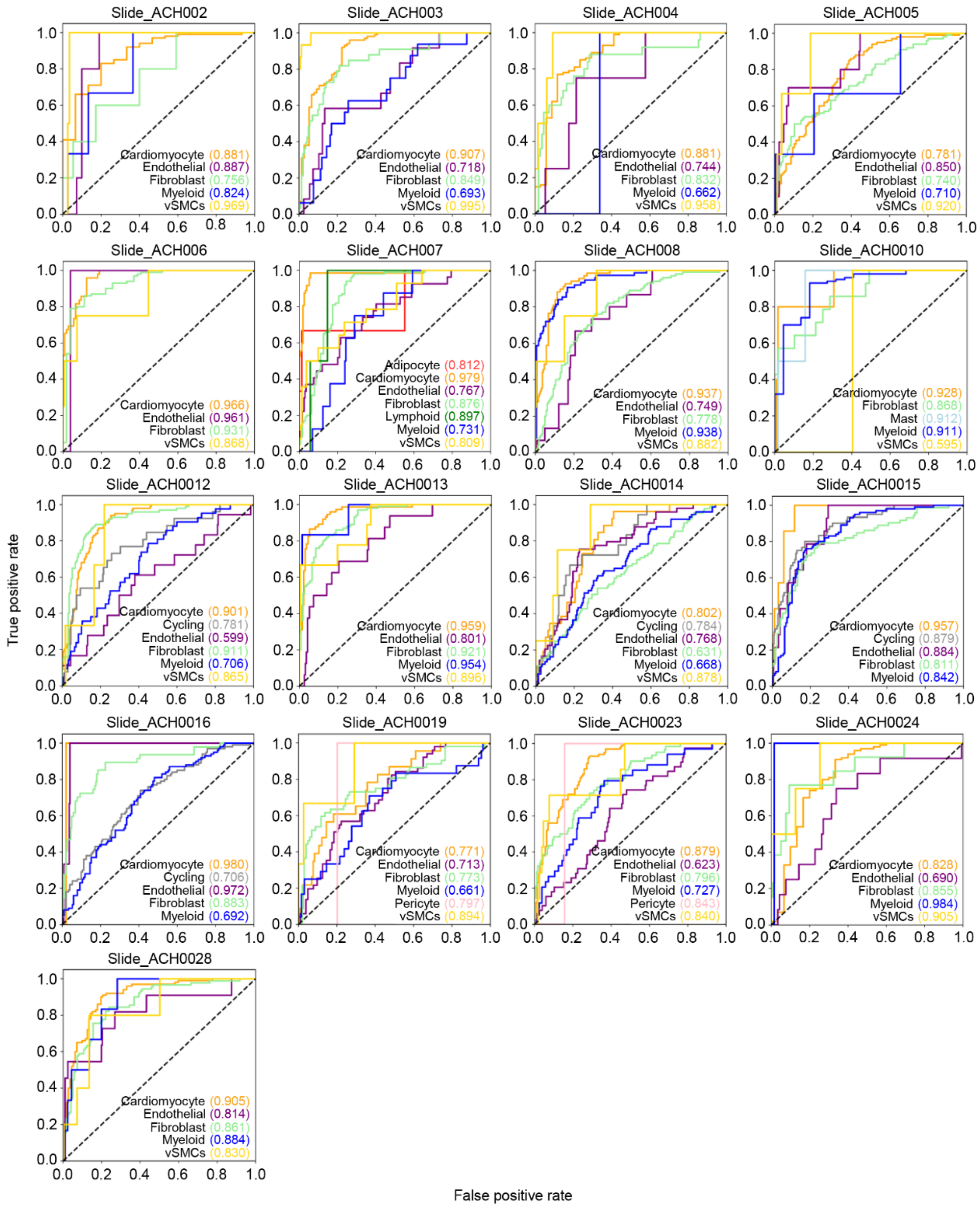


**Figure S10. Evaluation of STASCAN on the human cardiac dataset.** The ROC curves of each cell type by STASCAN on the human cardiac dataset. The number in each pair of parentheses represent the AUC value.

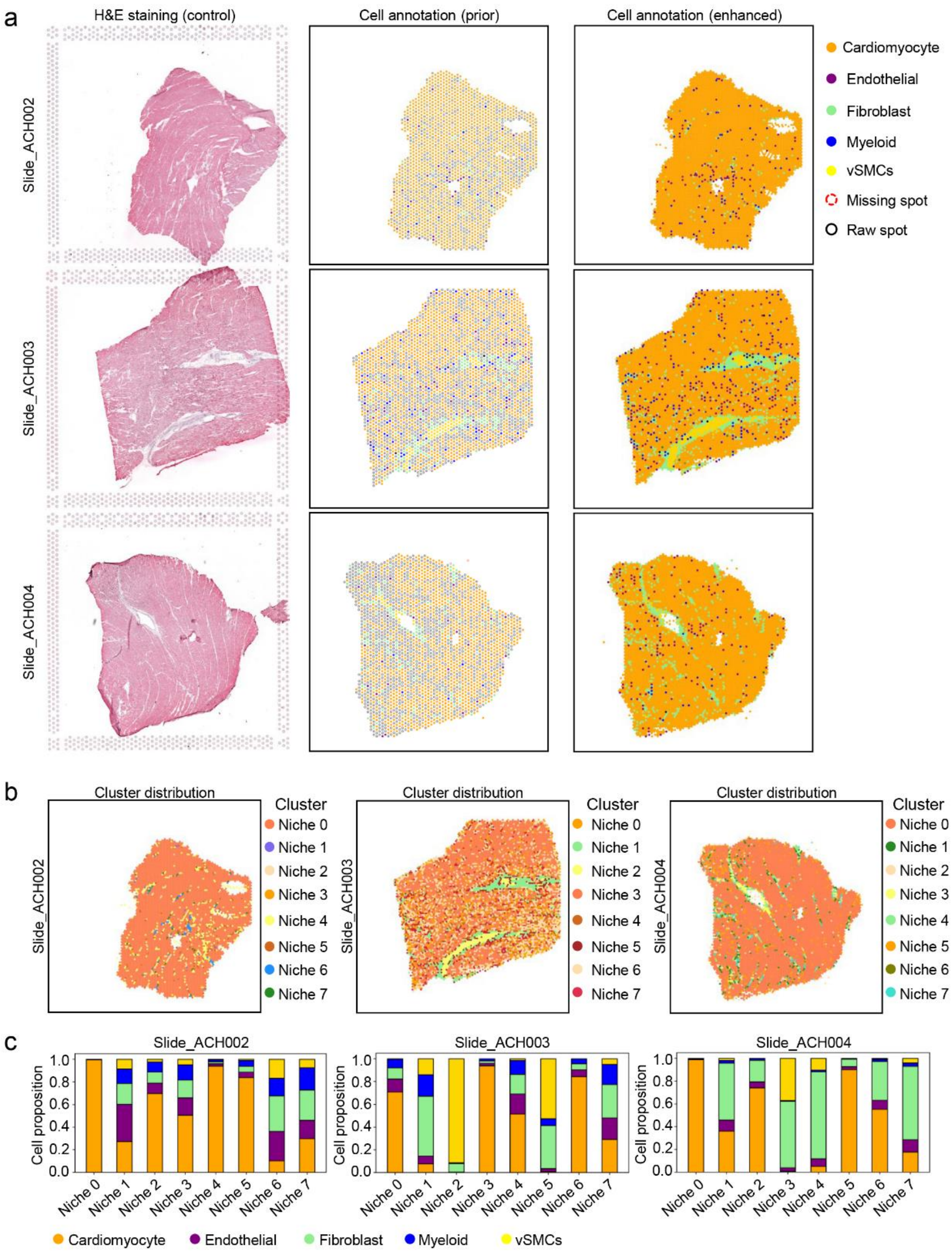


**Figure S11. Super-resolution cell distribution of slides sampled from non-transplanted donor hearts in the human cardiac dataset by STASCAN.**

**(a)** Tissue regions of the human cardiac from non-transplanted donor hearts (control). Left, H&E staining image showing general morphology. Middle, spatial plot (x and y axis) depicting cell distribution for the major cell types on prior spots. Right, spatial plot (x and y axis) depicting super-resolution cell distribution for the major cell types on the fully-tiled spots. Missing spots, spots removed from raw spots in the original literature due to the small number of measured genes and UMIs.

**(b)** Spatial plots (x and y axis) depicting spatial distribution for cell-type niches for each slide from non-transplanted donor hearts (control).

**(c)** Bar plot showing the cell proposition of cell-type niche for each slide from non-transplanted donor hearts (control).

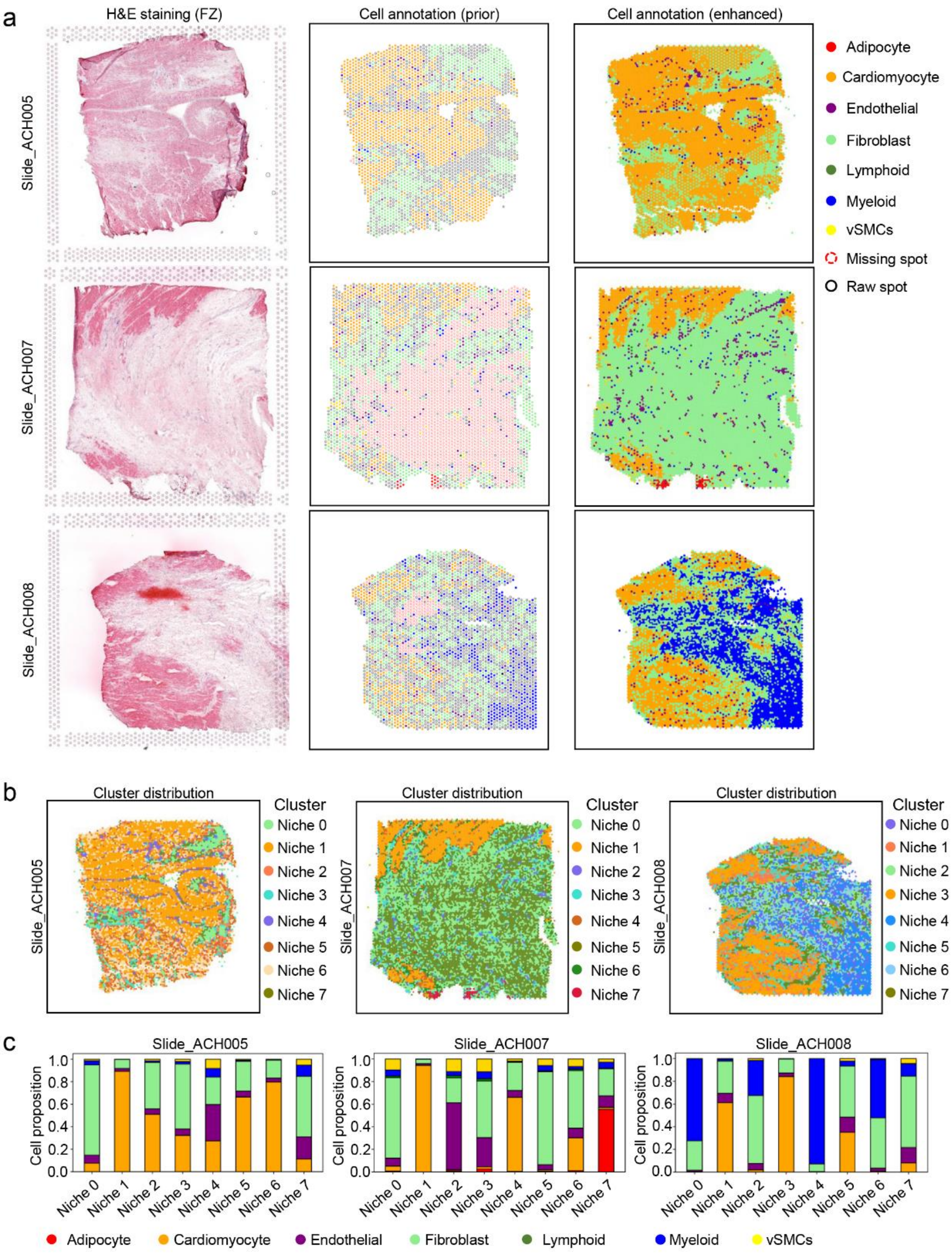


**Figure S12. Super-resolution cell distribution of slides sampled from the fibrotic zone in the human cardiac dataset by STASCAN.**

**(a)** Tissue regions of the human cardiac from the fibrotic zone, after myocardial infarction. Left, H&E staining image showing general morphology. Middle, spatial plot (x and y axis) depicting cell distribution for the major cell types on prior spots. Right, spatial plot (x and y axis) depicting super-resolution cell distribution for the major cell types on the fully-tiled spots. Missing spots, spots removed from raw spots in the original literature due to the small number of measured genes and UMIs. FZ, fibrotic zone.

**(b)** Spatial plots (x and y axis) depicting spatial distribution for cell-type niches for each slide from the fibrotic zone.

**(c)** Bar plot showing the cell proposition of cell-type niches for each slide from the fibrotic zone.

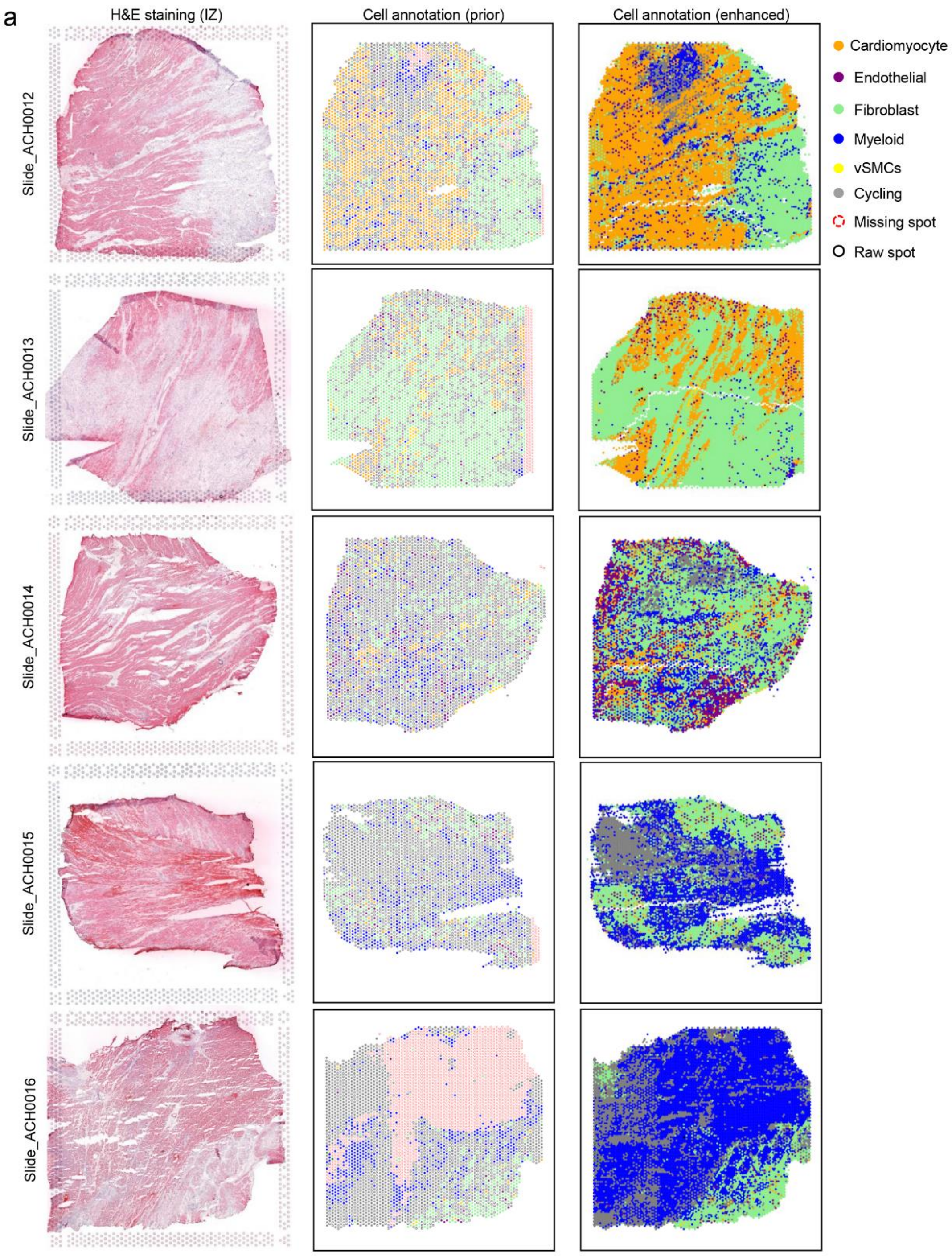
a
H&E staining (IZ)
Cell annotation (prior)
Cell annotation (enhanced)
Slide_ACH0012
Slide_ACH0013
Slide_ACH0014
Slide_ACH0015
Slide_ACH0016
Cardiomyocyte
Endothelial
Fibroblast
Myeloid
vSMCs
Cycling
Missing spot
Raw spot

**Next page**

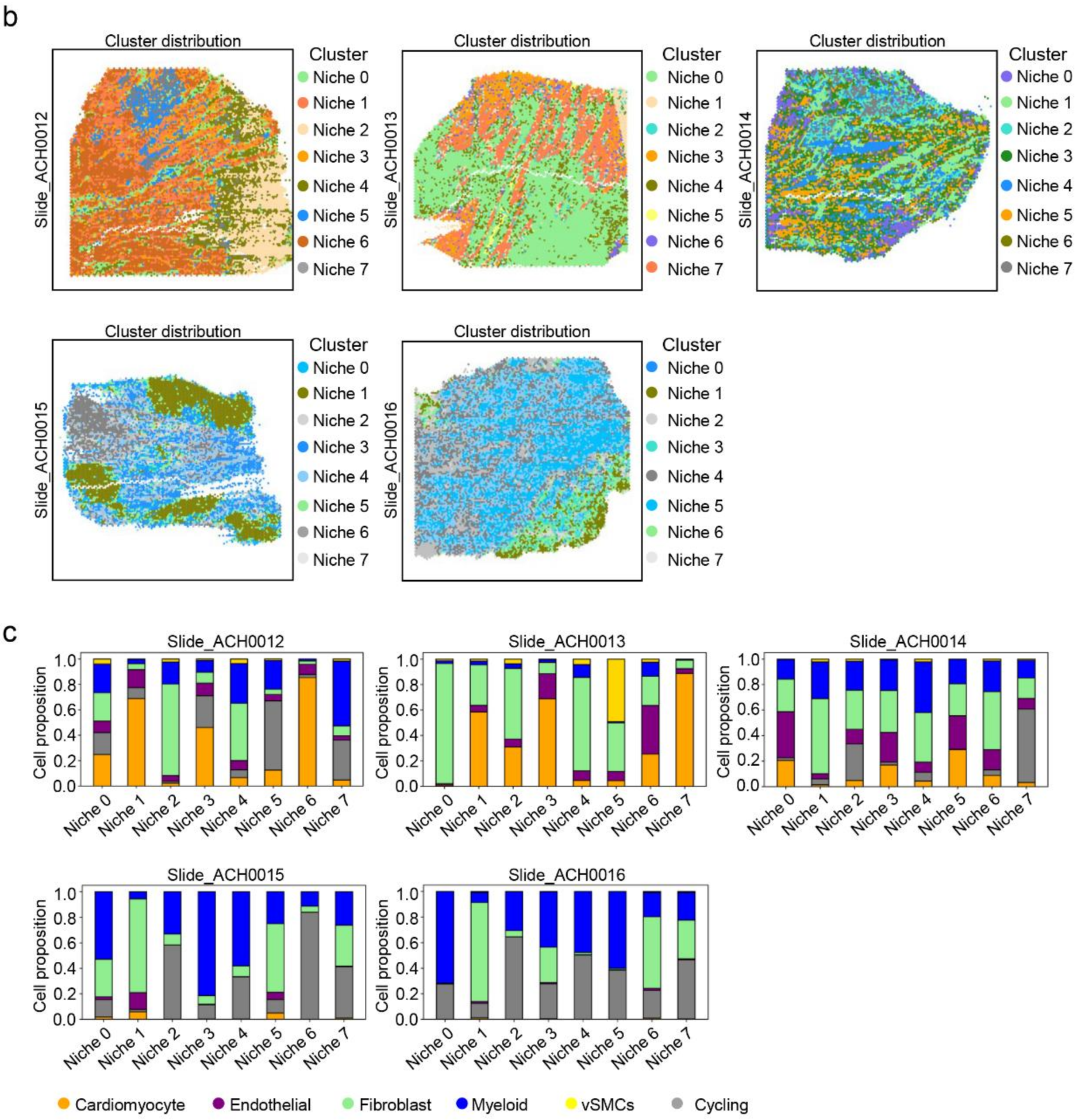


**Figure S13. Super-resolution cell distribution of slides sampled from the ischaemic zone in the human cardiac dataset by STASCAN.**

**(a)** Tissue regions of the human cardiac from the ischaemic zone, after myocardial infarction. Left, H&E staining image showing general morphology. Middle, spatial plot (x and y axis) depicting cell distribution for the major cell types on prior spots. Right, spatial plot (x and y axis) depicting super-resolution cell distribution for the major cell types on the fully-tiled spots. Missing spots, spots removed from raw spots in the original literature due to the small number of measured genes and UMIs. IZ, ischaemic zone.

**(b)** Spatial plots (x and y axis) depicting spatial distribution for cell-type niches for each slide from the ischaemic zone.

**(c)** Bar plot showing the cell proposition of cell-type niches for each slide from the ischaemic zone.

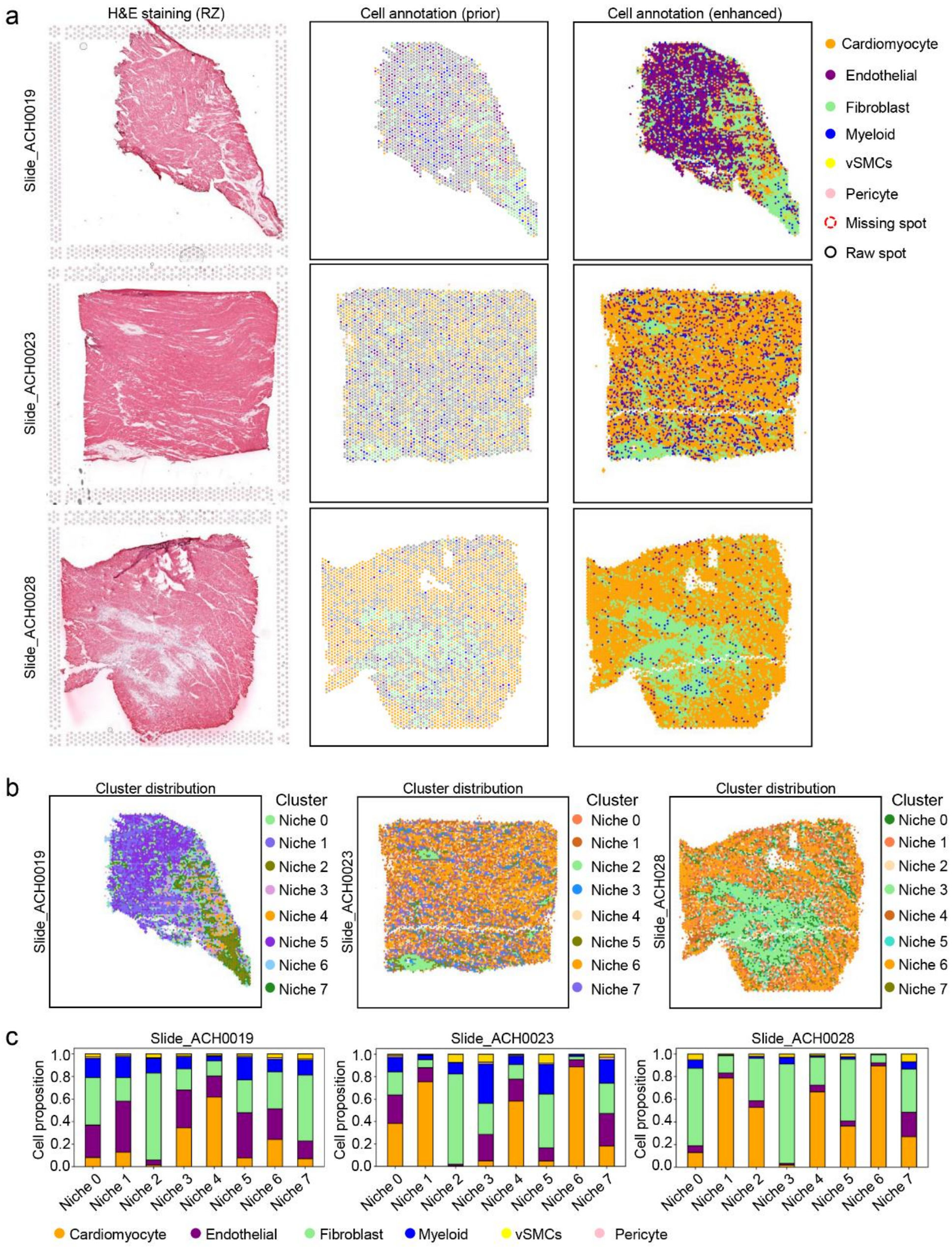


**Figure S14. Super-resolution cell distribution of slides sampled from the remote zone in the human cardiac dataset by STASCAN.**

**(a)** Tissue regions of the human cardiac from the remote zone, after myocardial infarction. Left, H&E staining image showing general morphology. Middle, spatial plot (x and y axis) depicting cell distribution for the major cell types on prior spots. Right, spatial plot (x and y axis) depicting super-resolution cell distribution for the major types on the fully-tiled spots. Missing spots, spots removed from raw spots in the original literature due to the small number of measured genes and UMIs. RZ, remote zone.

**(b)** Spatial plots (x and y axis) depicting spatial distribution for cell-type niches for each slide from the remote zone.

**(c)** Bar plot showing the cell proposition of cell-type niches for each slide from the remote zone.

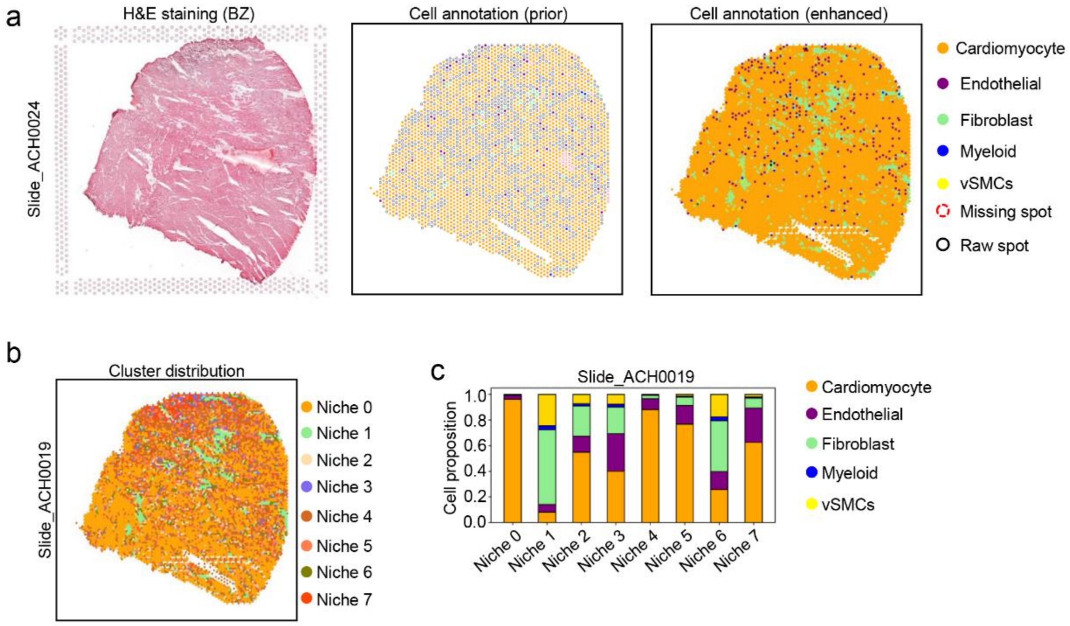


**Figure S15. Super-resolution cell distribution of the slide sampled from the border zone in the human cardiac dataset by STASCAN.**

(a) Tissue regions of the human cardiac from border zone, after myocardial infarction. Middle, spatial plot (x and y axis) depicting cell distribution for the major cell types on prior spots. Right, spatial plot (x and y axis) depicting super-resolution cell distribution for the major types on the fully-tiled spots. Missing spots, spots removed from raw spots in the original literature due to the small number of measured genes and UMIs. BZ, border zone.

(b) Spatial plots (x and y axis) depicting spatial distribution for cell-type niches from the border zone.

(c) Bar plot showing the cell proposition of cell-type niches from the border zone.
